# Genome-wide discovery of *cis-*regulatory elements in a large genome

**DOI:** 10.64898/2026.03.07.709930

**Authors:** Gillian Forbes, Emilia Skafida, Irene Karapidaki, Savannah Moinet, Mowgli Dandamudi, Çağrı Çevrim, Farzaneh Momtazi, Chryssa Anastasiadou, Sabrina Lo Brutto, Michalis Averof, Mathilde Paris

## Abstract

Identifying non-coding regulatory elements in the genome poses a challenge in most organisms. Classical methods rely on trial and error to test the regulatory activities of DNA fragments using reporter constructs. In large eukaryotic genomes, where *cis-*regulatory elements can spread over long distances, separated by large stretches of non-functional DNA, this trial and error approach is particularly challenging. Here, we generate two types of resources that can be used to narrow the search for such *cis-*regulatory elements in the 3.6 Gbp genome of *Parhyale hawaiensis* (comparable in size to the human genome). First, we use bulk ATAC-seq to uncover genome-wide patterns of chromatin accessibility in embryonic and adult tissues of *Parhyale* (whole embryos and legs), and single-nucleus ATAC-seq to identify regions of open chromatin in diverse cell types recovered from adult legs, including epidermal, neuronal, muscle and blood cells. Second, by sequencing the genomes of three congeneric species of *Parhyale hawaiensis* – *P. darvishi*, *P. aquilina* and *P. plumicornis* – we identify islands of sequence conservation across the genome, corresponding to DNA elements that are functionally constrained during evolution. We present an approach by which low-coverage (10-15x) short-read genome sequencing, without genome assembly, is sufficient to provide reliable maps of sequence conservation. This approach cuts the cost and labour required to generate these maps, making the identification of *cis-*regulatory elements more widely accessible. We demonstrate the utility of these resources by identifying *cis-*regulatory elements that drive robust expression of fluorescent reporters ubiquitously and in specific cell types.

## Introduction

Identifying *cis-*regulatory elements (CREs) that regulate transcription is key for understanding how gene expression is regulated in space and time, and for targeting the expression of genetic markers and tools to specific populations of cells. The classic approach for identifying CREs is based on trial and error: DNA fragments flanking the gene of interest are selected and their activity is tested using reporter constructs (e.g. Goto et al. 1989; Arnold et al. 2013; Pfeiffer et al. 2008). In organisms with compact genomes, CREs can often be found in proximity to promoters. In large eukaryotic genomes, however, CREs can be located tens or even hundreds of kilobases away from coding sequences (e.g. Sanyal et al. 2012; Levo et al. 2022). In the latter case, the task of identifying active *cis* elements within a sea of non-functional DNA is laborious and inefficient. Large research communities working in established model organisms (e.g. flies and mice) or humans, have historically invested substantial resources in identifying diverse CREs to support tool development (e.g. Arnold et al. 2013; Pfeiffer et al. 2008), but smaller communities working on non-conventional model organisms usually lack these resources (e.g. Sun et al. 2022; Lai et al. 2018).

A number of methods can be used to narrow down the candidate fragments of DNA to be tested. One approach is based on chromatin profiling. Transcriptional enhancers tend to be located in regions of ‘open’ chromatin, allowing access to transcription factors, they are often enriched in specific histone marks (e.g. H3K4me1 and H3K27ac) and maintain contacts with their target promoters. These features can be revealed by experimental techniques such as ATAC-seq, ChIP-seq and Hi-C, respectively (Buenrostro et al. 2013; Lieberman-Aiden et al. 2009; Barski et al. 2007). Given that each genomic locus is found in just two copies per diploid nucleus, a major constraint in using these methods is the number of cells required to obtain a robust signal. ChIP-seq and Hi-C typically require tens of thousands to millions of nuclei per experiment (Kidder et al. 2011; Belton et al. 2012). ATAC-seq requires lower amounts of starting material – hundreds to tens of thousands of nuclei per experiment (Buenrostro et al. 2013) – and has recently even been used in the context of single-cell profiling (Buenrostro et al. 2015). Chromatin accessibility profiles can therefore be obtained in the majority of experimental systems via ATAC-seq.

A second approach for identifying candidate CREs is to look for islands of DNA that show a high degree of sequence conservation during evolution, an approach known as phylogenetic footprinting (e.g. Duret and Bucher 1997; Blanchette and Tompa 2002; Pennacchio et al. 2006; Woolfe et al. 2005). The slower rate of evolution, resulting from functional constraints, helps to distinguish CREs from surrounding non-coding sequences that have no function. Comparisons between species that are separated by tens (and sometimes hundreds) of millions of years of evolution are effective in differentiating putative CREs from their surrounding non-functional sequences. Sequence conservation is often lost entirely over longer evolutionary timescales. To perform these comparisons, species at the appropriate evolutionary distances must therefore be collected and sequenced. A common limitation of this approach is the labour and cost involved in genome sequencing and assembly, particularly in species with very large genomes. We propose a workaround to this problem in this paper.

A third approach for identifying putative CREs has emerged recently with the development of deep learning models capable of predicting enhancer sequences (de Almeida et al. 2022; Barbadilla-Martínez et al. 2025). To generate reliable predictions, these models must be trained on large experimental datasets, which have so far been available only for *Drosophila* and mammals. It is not yet clear whether these models can perform well in other, distant species.

Here, we present resources for the discovery of CREs in the crustacean *Parhyale hawaiensis*, an emerging experimental model in comparative developmental biology, regeneration, chronobiology and ecotoxicology (reviewed in Paris et al. 2022; Averof 2022). *Parhyale* presents the typical challenges for CRE discovery in organisms with large genomes, having a genome of approximately 3.6 billion base pairs, similar in size to the human genome (Kao et al. 2016). Previous efforts to identify CREs relied on testing the activity of DNA fragments lying within a few kb from promoters (Pavlopoulos and Averof 2005; Pavlopoulos et al. 2009; Sun et al. 2022; Ramos et al. 2019). These efforts succeeded in identifying CREs from hsp70 and opsin genes, but failed to identify additional CREs, particularly ones associated with development or cell differentiation.

We combined two orthogonal approaches – chromatin profiling and sequence conservation – to identify putative CREs in the *Parhyale* genome. For this purpose, we produced the following resources: (1) bulk ATAC-seq data from embryonic and adult legs of *Parhyale*, (2) single-nucleus ATAC-seq data from adult legs of *Parhyale*, which yield the chromatin profiles of >15 cell clusters, including epidermis, muscles, neurons and blood, and (3) genome-wide maps of sequence conservation, highlighting islands of conservation in relation to other *Parhyale* species.

We show that it is not necessary to sequence genomes at high coverage, nor to generate genome assemblies, in order to identify islands of sequence conservation. Instead, we present a method that relies on mapping short sequence reads from new species (in this case, from *P. darvishi*, *P. aquilina* and *P. plumicornis*) with low stringency onto the genome assembly of the species of interest (*Parhyale hawaiensis*; Kao et al. 2016). This approach cuts the cost and labour required to generate such comparative data very significantly.

To demonstrate the utility of these resources, we have used them to discover both ubiquitous and cell-type specific CREs in the genome of *Parhyale*. First, we used the single-nucleus ATAC-seq data to identify regions of open chromatin that are shared by all cells, or regions that are specific to particular cell types such as neurons or muscles. Among these regions, we selected ones that show sequence conservation among related *Parhyale* species. Putative CREs were then tested using transgenic reporters, identifying two CREs that drive ubiquitous expression (out of 2 tested), two that drive expression specifically in neurons (out of 7 tested) and two that drive expression in muscles (out of 2 tested). Besides the value of these resources for the *Parhyale* research community, the labour- and cost-efficient approach we present here can be used to generate resources for CRE discovery in other species with large genomes.

## Results and Discussion

### Genome-wide profiles of chromatin accessibility in embryonic and adult tissues

To explore chromatin accessibility profiles across the *Parhyale* genome, we performed bulk ATAC-seq on nuclei isolated from three types of tissue: (i) single *Parhyale* embryos collected at stages S20, S23 and S24, covering embryonic stages when many organ systems are being formed (datasets E20, E23 and E24, respectively), (ii) developing legs collected at embryonic stages S25 to 26 (datasets EL1 to EL3), and (iii) fully differentiated adult T4 and T5 thoracic legs (datasets L1 to L3). These datasets identify tens of thousands of peaks of open chromatin distributed across 2.8 Gbp of the assembled *Parhyale* genome (results summarised in Table 1). These ATAC-seq data show a relatively low abundance of accessible chromatin in exons, introns and intergenic regions, and a strong enrichment of accessible chromatin surrounding transcription start sites (TSS; Figure 1B,D). The quality of ATAC-seq data is often measured by the fraction of reads that are found in peaks of accessible chromatin (FRiP); based on this criterion, our highest quality datasets are EL1 to EL3, L1 and E24 (see Table 1).

**Figure 1.**
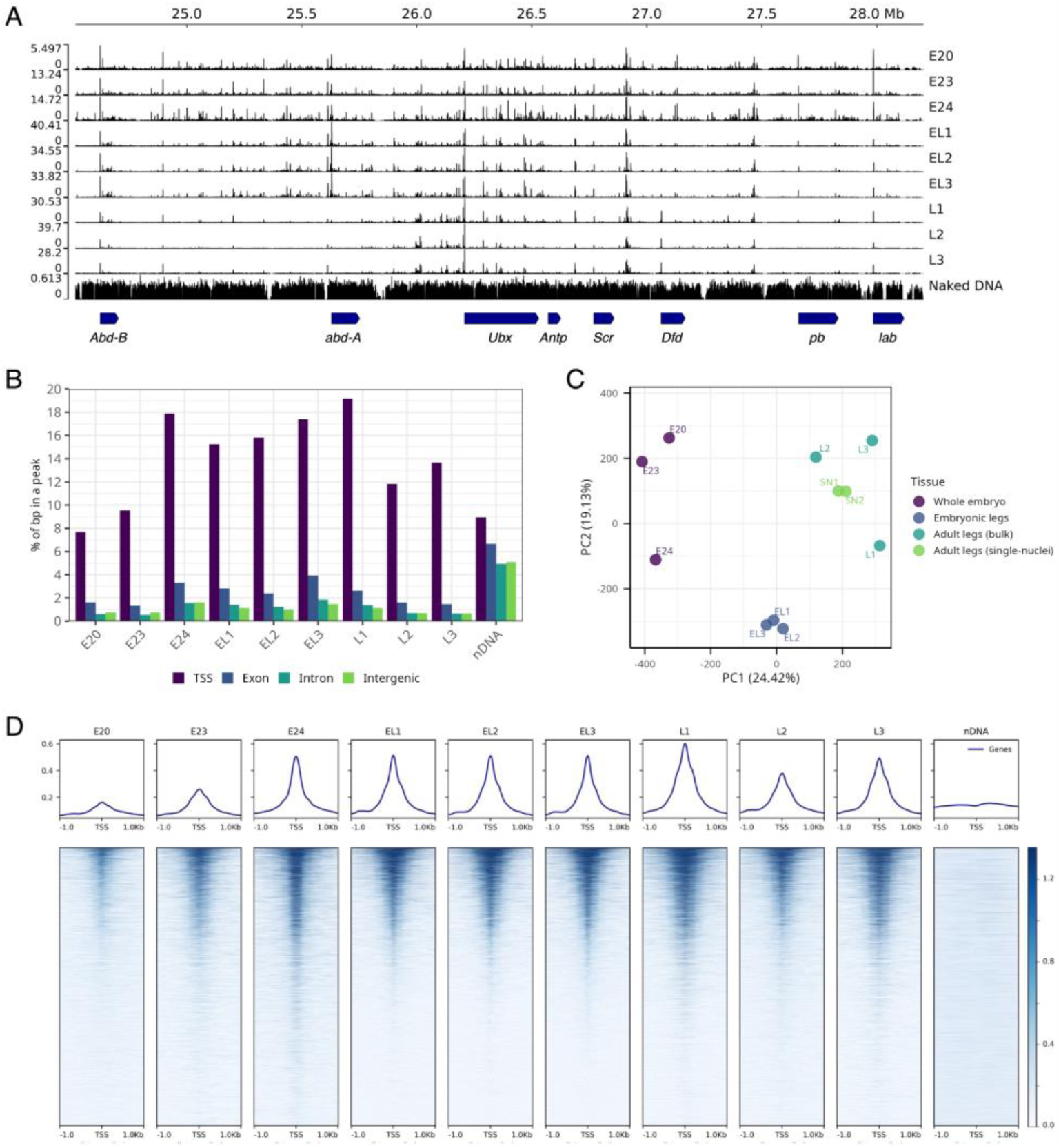
Comparison of ATAC-seq data from embryonic and adult tissues of *Parhyale hawaiensis*. (**A**) Genome browser plot of ATAC-seq data, focused on the 3.5 Mb spanning Hox gene cluster of *Parhyale hawaiensis*. In whole embryos (E20-24), ATAC-seq peaks are distributed across the entire Hox cluster. In embryonic thoracic legs (EL1-3), ATAC-seq peaks in the region of anterior and posterior Hox genes are partly suppressed. In adult T4 and T5 legs (L1-3), peaks of open chromatin are mostly visible in the region of *Ubx* and suppressed in other parts of the Hox cluster. (**B**) Histogram showing the proportion of transcription start sites (TSS), exonic, intronic and intergenic sequences in the *Parhyale* genome that overlap with an ATAC-seq peak in each dataset. Only TSS and exons of intron-containing genes were used in this analysis, to exclude poorly annotated transcripts. (**C**) Principal Component Analysis of the ATAC-seq datasets, based on the number of reads assigned to each peak. PC1 and PC2 (accounting together for 44% of the variation) show the datasets clustering according to tissue of origin (whole embryos, embryonic legs, adult legs, marked in different colours). (**D**) Mapping of ATAC-seq reads to the sequences surrounding transcription start sites across the genome; 1 kb upstream to 1 kb downstream of each TSS is depicted in each line (half of the 27,955 TSS are shown, keeping the same order per column; see Methods). Colours represent the number of mapped reads (see Methods). We observe a clear enrichment of open chromatin surrounding the TSS.

**Table 1.**
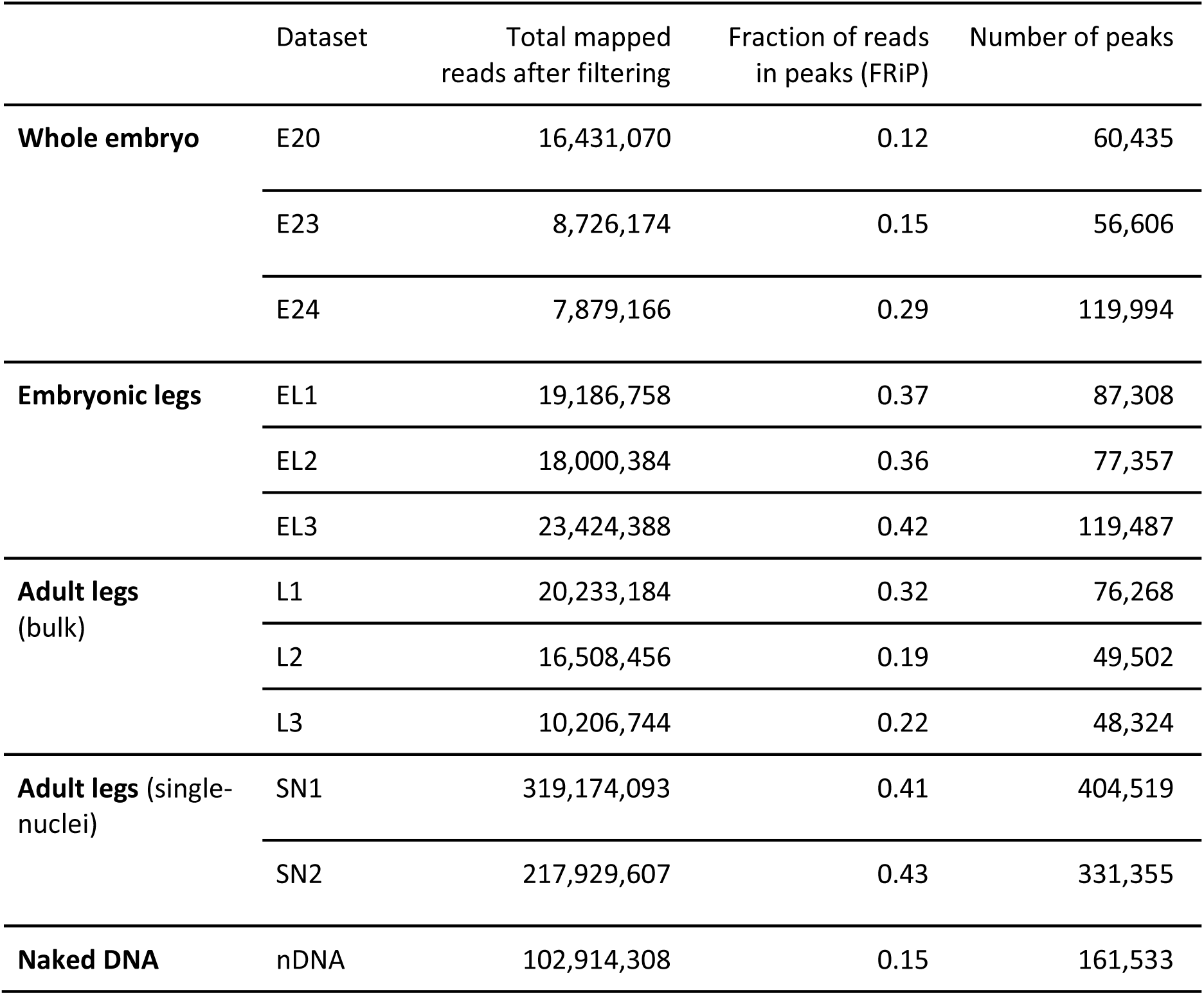
Overview of bulk and single-nuclei ATAC-seq datasets from *Parhyale hawaiensis*. Further information on mapping statistics are given in Suppl. Tables 1 and 2.

|  | Dataset | Total mapped reads after filtering | Fraction of reads in peaks (FRiP) | Number of peaks |
| --- | --- | --- | --- | --- |
| **Whole embryo** | E20 | 16,431,070 | 0.12 | 60,435 |
|  | E23 | 8,726,174 | 0.15 | 56,606 |
|  | E24 | 7,879,166 | 0.29 | 119,994 |
| **Embryonic legs** | EL1 | 19,186,758 | 0.37 | 87,308 |
|  | EL2 | 18,000,384 | 0.36 | 77,357 |
|  | EL3 | 23,424,388 | 0.42 | 119,487 |
| **Adult legs (bulk)** | L1 | 20,233,184 | 0.32 | 76,268 |
|  | L2 | 16,508,456 | 0.19 | 49,502 |
|  | L3 | 10,206,744 | 0.22 | 48,324 |
| **Adult legs (single-nuclei)** | SN1 | 319,174,093 | 0.41 | 404,519 |
|  | SN2 | 217,929,607 | 0.43 | 331,355 |
| **Naked DNA** | nDNA | 102,914,308 | 0.15 | 161,533 |

Principal Component Analysis, based on the number of reads assigned to each peak, shows these datasets clustering according to the origin of the samples (whole embryos, embryonic legs, adult legs), with the tightest clustering seen in the embryonic leg datasets (Figure 1C).

Some of the variation between samples is likely to reflect differences in chromatin accessibility between tissues and developmental stages. To test this, we focused on the genomic region that encompasses the *Parhyale* Hox genes. Based on the mechanisms of Hox gene regulation known to operate in *Drosophila* (Bowman et al. 2014; Peifer et al. 1987) and the expression of Hox genes in *Parhyale* (Serano et al. 2016), we expected open chromatin peaks to be distributed across the entire Hox cluster in embryos (E20, E23 and E24), but only the region surrounding the gene *Ubx* in adult T4 and T5 thoracic legs (L1-3). The patterns we observe partly match that expectation (Figure 1A).

### Genome-wide profiles of chromatin accessibility from diverse cell types

Many research projects on development and regeneration focus on gene regulation and expression profiles that are specific to particular cell types. We have previously shown that *Parhyale* legs contain more than 15 transcriptionally-distinct cell types, including epidermal and neuronal cells, muscle blood, and several yet unidentified cell types (Almazán et al. 2022). To probe the cellular specificity of chromatin landscapes, we performed single-nucleus ATAC-seq (snATAC-seq) on adult *Parhyale* T4 and T5 legs (datasets SN1 and SN2, comprising a total of 15,969 nuclei).

By pooling the ATAC-seq reads from all the cells captured in these datasets, we generated ‘pseudo-bulk’ SN1 and SN2 datasets which are comparable with the bulk ATAC-seq datasets (Table 1, Figures 1A,B). Principal Component Analysis shows that these datasets cluster with the bulk ATAC-seq datasets generated from adult legs (Figures 1C).

The snATAC-seq data identify clusters of cells with distinct chromatin landscapes (Figure 2A,B). Based on the ATAC-seq peaks surrounding cell-type-specific marker genes identified by snRNA-seq (Almazán et al. 2022), we are able to identify some of these cell clusters as epidermal, muscle, neuronal and blood cells (Figure 2C,D; see Methods). The cells of each cluster share ATAC-seq peaks that are specific to the cluster (e.g. Figure 2E). Thus, our snATAC-seq data identify genomic sequences that are differentially accessed in different cell types.

**Figure 2.**
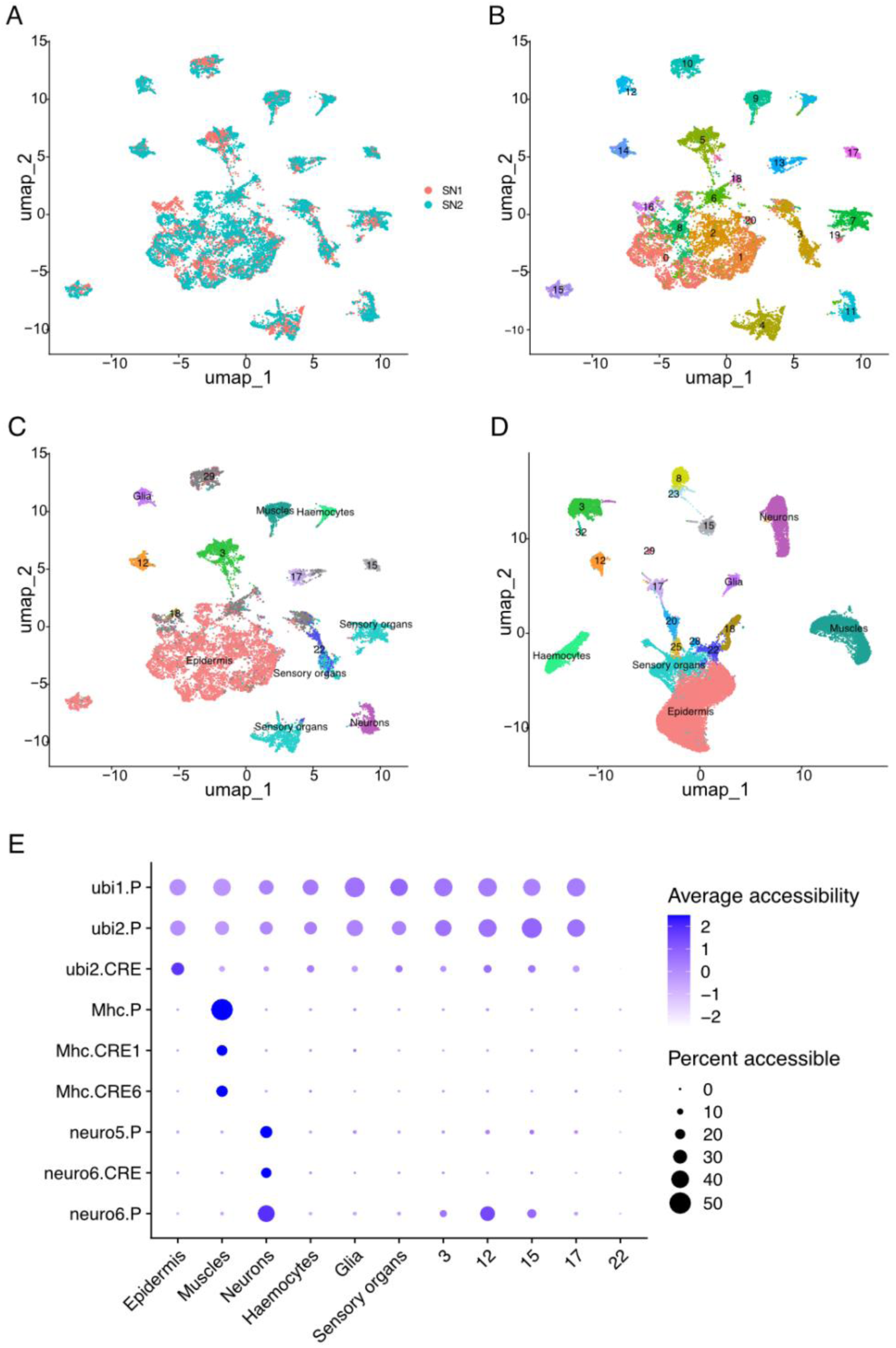
Single nuclei ATAC-seq and differential chromatin accessibility across cell types. (**A-C**) UMAP of integrated snATAC-seq experiments SN1 and SN2, including 15,969 cells from adult uninjured *Parhyale* legs (7,951 cells from SN1 and 8,018 cells from SN2). UMAP was colour-coded based on experiment (A), cell clusters identified from the ATAC-seq signal (B), or cell types identified from previously published snRNA-seq data (Almazán et al. 2022; see Methods, Suppl. Figure 2) (C). (**D**) UMAP of snRNA-seq data published in (Almazán et al. 2022), using the same colour code. (**E**) Dot plots of the ATAC-seq signal at the putative CREs that we tested *in vivo*, per cell cluster.

### *Parhyale* species available for comparative genomics

Our initial efforts to identify regions of sequence conservation in the *Parhyale* genome focused on comparisons with *Hyalella azteca*, a hyalid amphipod whose genome was sequenced in 2018 (Poynton et al. 2018). Comparing non-coding sequences between *Pahyale* and *Hyalella*, however, failed to reveal islands of sequence conservation, even in loci known to harbor functional CREs (Suppl. Figure 1). Phylogeographic studies estimate that the lineages of *Parhyale* and *Hyalella* likely diverged 110 to 140 million years ago (Cannizzaro and Berg 2022), likely too long for detecting sequence conservation in CREs.

To identify regions of sequence conservation over shorter evolutionary distances, we collected three species that belong to the same genus as *Parhyale hawaiensis* – *P. aquilina*, *P. darvishi* and *P. plumicornis* – and we sequenced their genomes to an estimated genome coverage of 16x, 11x and 10x, respectively (Table 2, Suppl. Figure 3; these values may represent slight overestimates, see Methods). Based on our analysis, the genome sizes of *P. aquilina*, *P. darvishi* and *P. plumicornis* are estimated to be 1.1 Gbp, 2.8 Gbp and 3.0 Gbp, respectively.

**Table 2.**
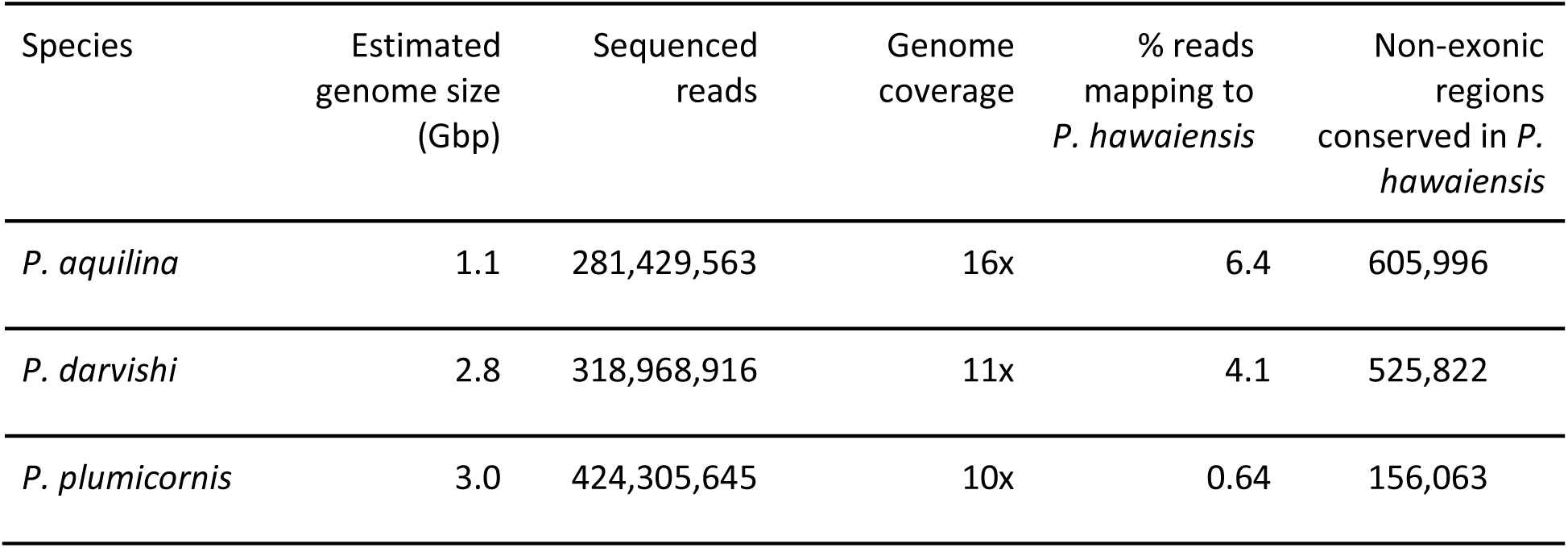
Overview of short-read genome sequencing on three *Parhyale* species.

The reads from each species were then mapped with low stringency to a genome assembly of *P. hawaiensis* (Phaw_5.0, GenBank GCA_001587735.2; updated in Almazán et al. 2022). Using this approach, 6.4% of reads from *P. aquilina*, 4.1% of reads from *P. darvishi*, and 0.46 % of reads from *P. plumicornis* could be mapped unambiguously to the *P. hawaiensis* genome (Table 2), suggesting that sequence comparisons within the genus *Parhyale* could be informative for CRE discovery in these animals. These levels of cross-species mapping likely reflect the relative phylogenetic proximity and genome sizes of these species. A molecular phylogeny based on the sequences of conserved single-copy genes (BUSCO gene set, 29,097 aligned nucleotides across all species; see Methods) reveals the phylogenetic relationships of these species (Figure 3A).

**Figure 3.**
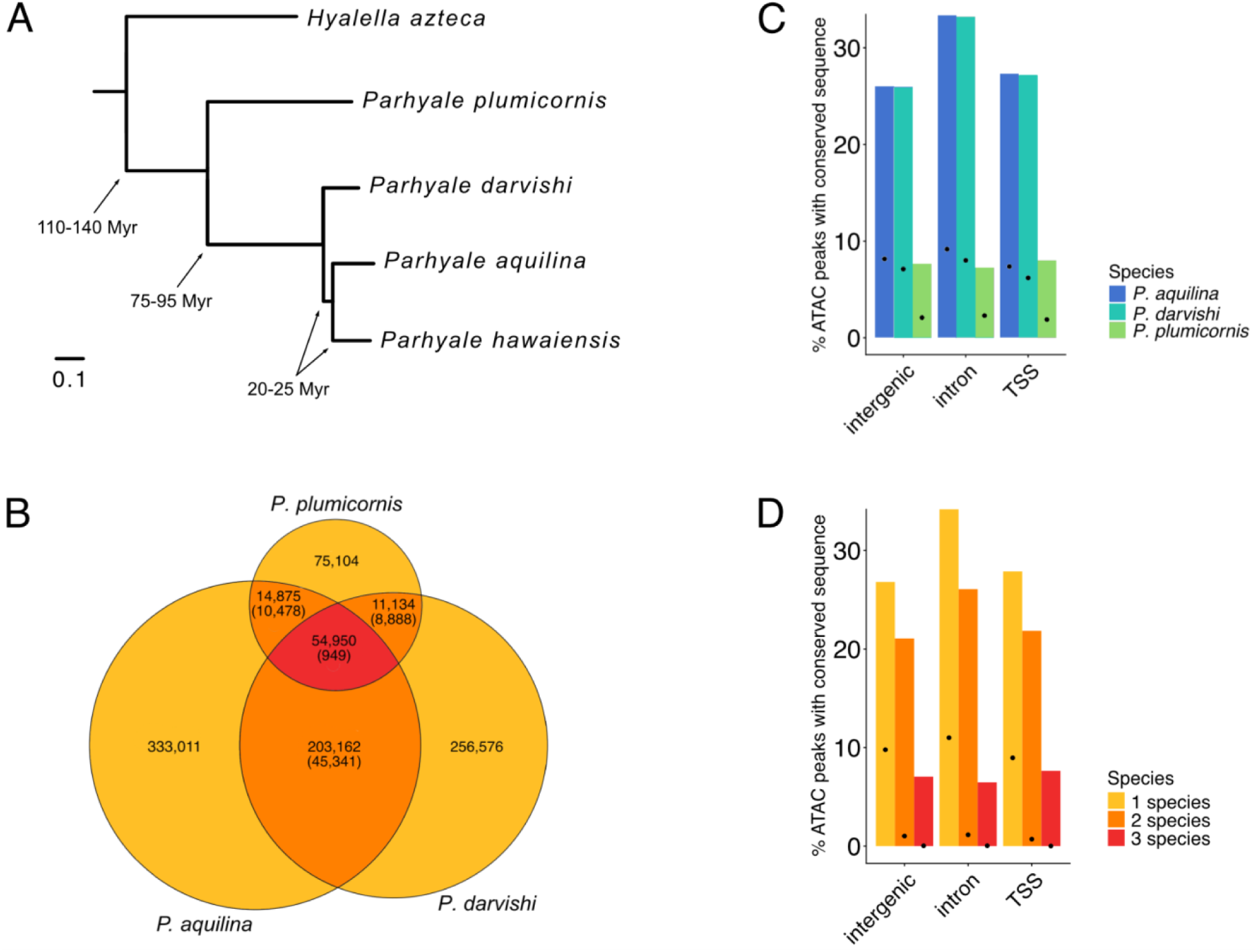
Cross-species sequence conservation and its relation with chromatin. (**A**) Molecular phylogeny depicting the evolutionary relationships of the *Parhyale* species included in this study (*P. hawaiensis*, *P. aquilina*, *P. darvishi* and *P. plumicornis*), with *Hyalella azteca* as the outgroup. The phylogeny was based on 29,097 aligned nucleotides from conserved single-copy genes (BUSCO gene set; see Methods). Divergence estimates (in million years) were calibrated against the estimated divergence between *Parhyale* and *Hyallella* (Cannizzaro and Berg 2022). Scale bar represents number of substitutions per site. (**B**) Overlap between regions of sequence conservation in non-exonic sequences (TSS, introns and intergenic regions) identified by mapping of reads from *P. aquilina*, *P. darvishi* and *P. plumicornis* onto the genome the *P. hawaiensis* (see Methods). Values in parentheses indicate the number of overlapping regions that would be expected if the conserved regions were distributed randomly. (**C**) Quantification of the fraction of ATAC-seq peaks in non-exonic sequences that overlap a conserved region identified in *P. aquilina*, *P. darvishi* and *P. plumicornis* (see Methods). (**D**) Quantification of the fraction of ATAC-seq peaks in non-exonic sequences that overlap a conserved region in either one, two or three of the other *Parhyale* species (see Methods). Colour code corresponds to panel B. Black dots in panels C and D represent the overlap that would be expected if the conserved regions were distributed randomly.

An estimate of divergence times based on the relative branch lengths in this tree, calibrated against *Hyalella azteca* (Cannizzaro and Berg 2022), suggests that *P. hawaiensis* has likely diverged for 20 to 25 million years from *P. aquilina* and *P. darvishi*, and 75 to 95 million years from *P. plumicornis*.

### Genome-wide patterns of sequence conservation

The cross-species read mapping identifies hundreds of thousands of islands of sequence conservation in the genome of *P. hawaiensis* (Table 2, Figure 3B). Of these, 8% are found in exons, likely reflecting conservation in protein coding sequences, while 37% are found in introns, 54% in intergenic regions, and 1% overlap with promoters (TSS), marking regions that evolve at a lower rate than surrounding non-coding sequences. The peaks in non-exonic sequences include 54,950 that show conservation across all four species. If these peaks of sequence conservation were distributed randomly in the genome, only 949 would be expected to overlap between all four species (Figure 3B). This suggests that a large proportion of these sequences is conserved due to constraints that are maintained over a scale of 100 million years of divergent evolution.

The islands of sequence conservation found in non-coding sequences (promoter/TSS regions, introns and intergenic regions) coincide with peaks of open chromatin identified by ATAC-seq more frequently than would be expected by chance (Figure 3C,D), consistent with the hypothesis that many of these regions represent *cis-*regulatory elements. The binding of regulatory proteins and RNA to these regions likely explains their chromatin accessibility and constrains their rate of sequence evolution.

### Identification of *cis-*regulatory elements driving ubiquitous, neuron- and muscle-specific expression

In the past, most attempts to identify ubiquitous or cell-type specific CREs in *Parhyale* using DNA fragments encompassing putative promoters, were unsuccessful. For example, in a recent effort to identify neuron-specific CREs, we generated reporter constructs carrying putative promoter regions from genes expressed in neurons (*bruchpilot*, *VAChT*, *synaptotagmin 4*) or involved in neuronal differentiation (*achaete/scute*, *senseless*, *cut*, *prospero*, *neuralised*). The 10 fragments we tested extended 1.5 to 5.5 kb upstream of the start codon, including a 3’UTR, putative TSS, and 1 to 5 kb of upstream sequences (see Methods). None of these reporters was found to be expressed in embryos. In retrospect, it appears that some of these fragments lacked the ATAC-seq peak that is usually associated with the TSS, suggesting that the promoter region had been incorrectly identified in the absence of ATAC-seq data.

To test the utility of the ATAC-seq and sequence conservation data we generated, we used these resources to identify CREs with ubiquitous and cell-type-specific activities. To identify ubiquitous promoters, we searched for genes that are robustly expressed in all cell clusters in adult *Parhyale* legs (Almazán et al. 2022); we then identified those with promoters that exhibit a clear ATAC-seq peak in all cell types, and overlap with sequences that are conserved in at least one other *Parhyale* species. Based on this approach, we identified two promoter regions and an associated putative CRE belonging to the *Parhyale* orthogues of the genes *headcase (hdc*, referred to as ubi1) and *muscleblind (mbl*, referred to as ubi2) (Figures 2E and 4A,B, and Table 3). To test the activity of these sequences, we generated reporter constructs carrying these fragments upstream of the coding sequence of mNeonGreen, and cloned these into the *Minos* transgenesis vector. We microinjected these constructs in early *Parhyale* embryos and we screened the resulting transgenic mosaic embryos by fluorescence microscopy (see Methods). We found that both *ubi1* and *ubi2* give strong widespread expression in at least 30% of the embryos and hatchlings we screened (Figure 4A’,A’’,B’,B’’).

**Figure 4.**
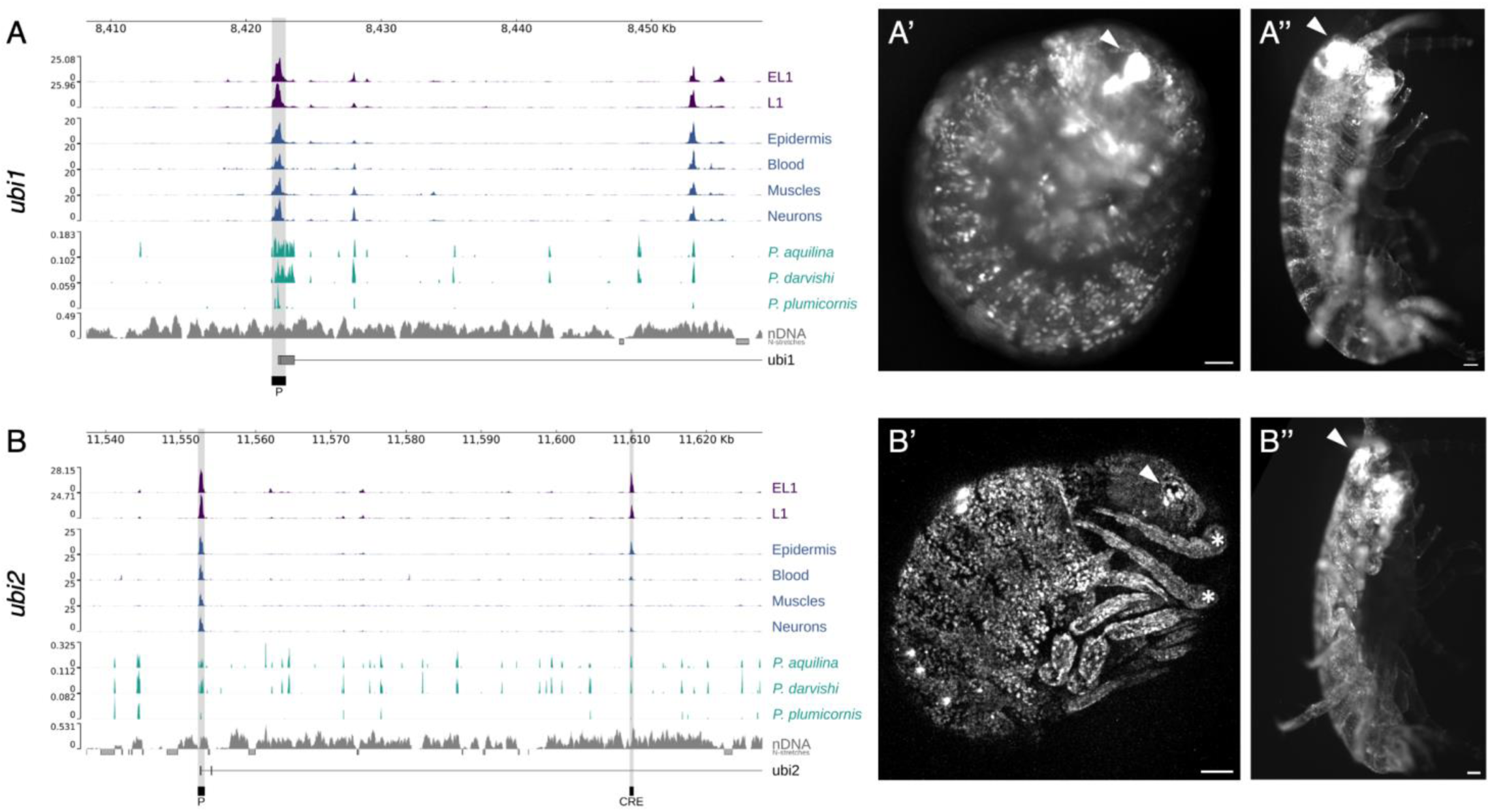
Identification of CREs driving ubiquitous expression. (**A,B**) Genome tracks showing the ATAC-seq and sequence conservation profiles that we used to identify the *ubi1* and *ubi2* promoters and associated CREs. (**A’,B’**) Fluorescence observed with the *ubi1.P* (A’) and *ubi2.CRE+P* (B’) reporters in live late stage embryos. Image A’ was captured on a fluorescence stereoscope, image B’ is a max projection of an image stack captured by confocal microscopy. Side views of embryos, with the head located at the top right, dorsal side towards the top-left (see Browne et al. 2005). Arrowheads mark the eye; asterisks in B’ mark the base of the antennae. (**A’’,B’’**) Fluorescence observed with the *ubi1.P* (A’’) and *ubi2.CRE+P* (B’’) reporters in live hatchlings. These images show ventro-lateral views of unilateral genetic mosaics, with expression only on one half of the animal captured on a fluorescence stereoscope. The head is located at the top, dorsal side on the left. Arrowheads mark the eye. Scale bars, 50 µm. The raw image data are available in Suppl. Data 8.

**Table 3.**
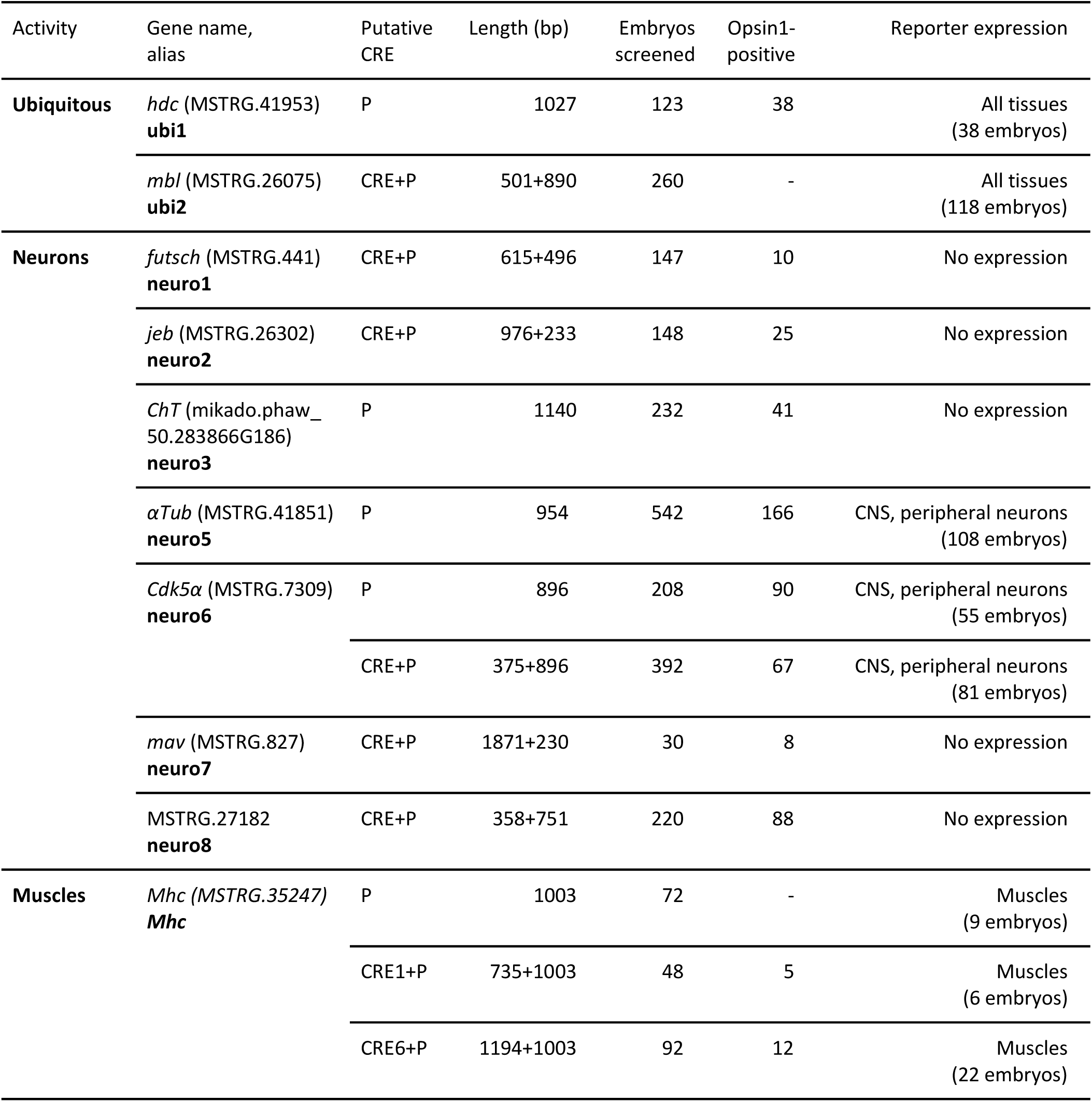
Putative CREs with ubiquitous, neuron- and muscle-specific activity. The transgenic reporters that we tested carry the promoter region (P) of the gene of interest and additional putative CREs upstream of the EGFP or mNeonGreen coding sequences. The fragments tested are shown in Figures 4A-B, 5A-C. The number of embryos screened, the number of embryos showing unilateral or bilateral expression of the Opsin1 transgenesis marker, and the number of embryos showing (mosaic) reporter expression are indicated. Genetic mosaicism results in different numbers of positives for the Opsin1 marker (eyes) and reporter expression (other tissues). CNS: central nervous system. The P and CRE sequences are given in Suppl. Data 7.

To identify neuron-specific CREs, we used the snATAC-seq data to identify 37 regions of the genome that are preferentially accessible in neurons (relative to other cell types) and located near genes that are preferentially expressed in neurons based on snRNA-seq data (Almazán et al. 2022). Among these regions, we selected ones that show sequence conservation in at least one other *Parhyale* species. Based on this analysis, we selected candidate promoters and associated putative CREs from 7 genes (Table 3, Figure 5A,B, Suppl. Figure 4). To test their activities, we generated reporter constructs carrying these promoters and CREs upstream of the coding sequence of EGFP or mNeonGreen, cloned these into the *Minos* transgenesis vector (see Methods) and microinjected each construct in early *Parhyale* embryos. The resulting transgenic mosaics were screened at late embryonic stages. Two of the 7 reporters we tested, named *neuro5* and *neuro6* (carrying CREs associated with the genes *αTub* and *Cdk5α*, respectively), showed strong fluorescence in the brain, ventral nerve cord, and peripheral neurons in the body, legs and antennae (Figure 5A’,A’’,B’,B’’). We observed these patterns of fluorescence in approximately 20% of the surviving injected embryos. The *neuro6* promoter alone gave expression in the nervous system, suggesting that at least part of the *neuro6* reporter activity resides in the promoter. Transgenic animals carrying both *neuro5* and *neuro6* reporters show that these two CREs drive expression in distinct subsets of neurons (Suppl. Figure 5). The embryos injected with the other five constructs showed no significant fluorescence above background levels.

**Figure 5.**
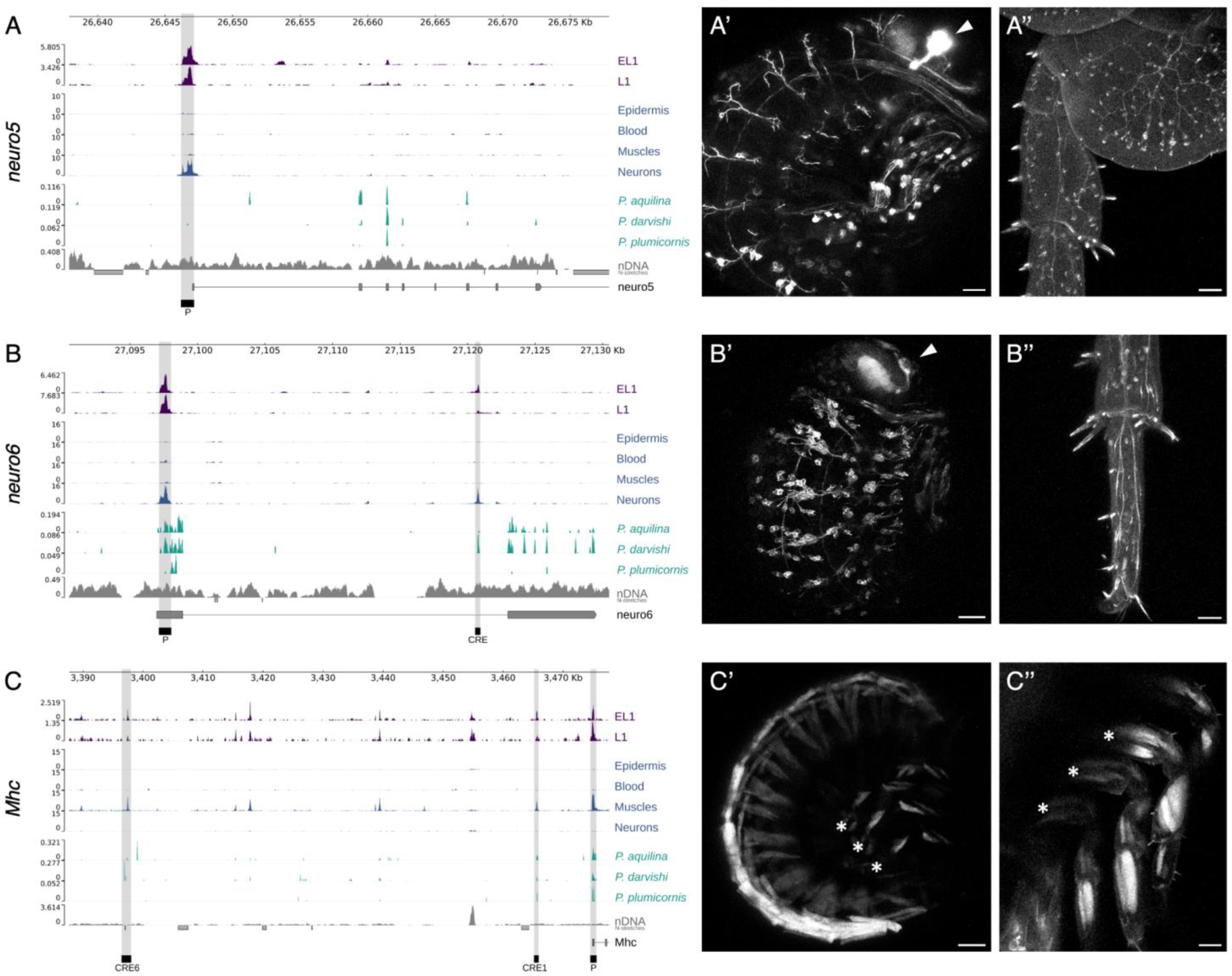
Identification of CREs driving expression in neurons and muscles. (**A-C**) Genome tracks showing the ATAC-seq and sequence conservation profiles that we used to identify the *neuro5*, *neuro6* and *Mhc* neuron- and muscle-specific promoters and associated CREs. (**A’-C’**) Fluorescence observed with the *neuro5-Src64B-mNeonGreen* (A’), *neuro6-Src64B-mNeonGreen* (B’) and *Mhc.CRE6+P-mNeonGreen* (C’) reporters in late embryos. Max projections of image stacks captured by confocal microscopy. Side views of embryos, with the head located at the top right, dorsal side towards the top-left. Arrowheads mark the eye; asterisks in C’ mark the base of thoracic legs. (**A’’-C’’**) Fluorescence observed with the of the *neuro5.P* (A’’), *neuro6.P+CRE* (B’’) and *Mhc.CRE6+P* (C’’) reporters in the thoracic legs of juveniles (proximal part of the leg including coxal plate in A’’, distal tip of leg in B’’, proximal part of several legs in C’’). All panels show max projections of image stacks captured by confocal microscopy. Asterisks in C’’ mark the base of thoracic legs. Scale bars, 50 µm. The raw image data are available in Suppl. Data 8.

To identify muscle-specific CREs, we focused on the genomic region surrounding the *Myosin heavy chain* gene (*Mhc*, MSTRG.35247), which is strongly and specifically expressed in nuclei of the muscle cluster of *Parhyale* (Almazán et al. 2022). We identified regions in which chromatin is specifically accessible in the muscle cluster and show sequence conservation across *Parhyale* species (Figure 5C), and tested the activity of two of these regions using the reporter approach described earlier. In both reporters, we observed specific expression in the muscles in 12-23% of the surviving injected embryos (Figure 5C’,C’’). Reporters with the Mhc promoter alone gave similar expression, suggesting that at least part of this activity resides in the promoter.

Evidence from diverse sources suggests there are different classes of *cis-*regulatory elements associated with different types of genes, namely with ubiquitously expressed ‘housekeeping’ genes, genes expressed in terminally differentiated cells, and developmentally-regulated genes. Each class tends to be associated with different sequence motifs and exhibits differences in nucleosome positioning, precision of transcription initiation and the use of distantly located enhancers (reviewed by Haberle and Lenhard 2016). To test whether our approach for identifying CREs can be effective in identifying the enhancers of developmental genes, we used the ATAC-seq data from embryonic legs (EL1-3) combined with cross-species sequence conservation, to identify putative enhancers in three leg patterning genes, *Dll-e*, *dac1* and *dac2* (Bruce and Patel 2020; Kao et al. 2016; Pavlopoulos et al. 2009). We tested the activity of 13 distant CREs using fluorescent reporter constructs, carrying each CRE with the promoter region of the corresponding gene (Suppl. Table 3 and Suppl. Figure 6). Among these constructs, we could only detect weak ubiquitous expression driven by promoter-proximal sequences of *Dll-e* (Suppl. Table 3). We did not find constructs that recapitulate the endogenous pattern of *Dll-e*, *dac1* and *dac2* expression in the legs of *Parhyale*. This result may reflect either our inability to identify the enhancers regulating these developmental genes, or expression levels that are below the detection threshold of our fluorescent reporters.

## Conclusions

Previous efforts to identify cis-regulatory elements in *Parhyale* relied on reporter constructs carrying a few kb of sequence upstream of selected target genes, an approach that has worked well in animals and plants with relatively small genomes. As presented earlier, however, this approach was often unsuccessful in *Parhyale*: from tens of reporters tested, only four robust native drivers had so far been identified (Pavlopoulos and Averof 2005; Pavlopoulos et al. 2009; Sun et al. 2022; Ramos et al. 2019). The present work adds 5 new drivers to that collection, including ones with ubiquitous, neuron- and muscle-specific activities. This result comes from combining information on genome-wide chromatin accessibility and evolutionary conservation profiles.

We cannot at this point distinguish the relative contributions of chromatin profiling and sequence conservation to CRE discovery, because these sources of information were not tested separately. At minimum, we can state that (1) ATAC-seq profiles serve to identify robustly the promoters of candidate genes that are active in a particular cellular context (cell type and stage) and (2) coupling this information with sequence conservation narrows down candidate promoters and distant CREs by a factor of 4 to 10, since only a fraction of ATAC-seq peaks show sequence conservation (Figure 3D). This represents a great improvement in our ability to select candidates to test by transgenesis, the most labour intensive step in the process.

Our approach relies on ATAC-seq and low-coverage short-read genome sequencing, which are today widely accessible and affordable. Both methods can be applied to small samples: a few thousand cells can be sufficient for ATAC-seq (as in our single-embryo samples) and a few micrograms of DNA are sufficient for low-coverage short read sequencing of most genomes. These methods can be combined with existing genome annotations and information on gene expression or function, to select relevant candidate genes. There are therefore few barriers to adopting this approach.

For ATAC-seq, good quality nuclear preparations are necessary, which could limit the adoption of this method in cases where cells are embedded within a tough extracellular matrix or a highly resistant cell wall. To detect sequence conservation by low coverage genome sequencing, the phylogenetic distance between species is a key factor. The approach is most powerful when evolution has had sufficient time to erase sequence similarities in non-functional (neutrally evolving) sequences, but not to eliminate conservation within functionally constrained sequences. This degree of sequence divergence is usually found in species separated by tens of millions of years of evolution. In *Parhyale*, we have found that sequence comparisons between *P. hawaiensis*, *P. aquilina*, *P. darvishi* and to some extent *P. plumicornis* reveal islands of sequence conservation in non-exonic sequences that cannot be found in comparisons with more distant relatives such as *Hyalella* (e.g. see Suppl. Figures 1 and 7).

Our strategy of mapping regions of sequence conservation by direct mapping of short sequence reads across species is much more accessible than conventional strategies that rely on genome assembly. The latter require much higher sequence coverage (> 50x) from multiple libraries, long-read sequencing or other scaffolding methods, and complex bioinformatic pipelines to assemble large genomes. Moreover, these approaches are often compromised by high levels of polymorphism found in natural populations. We estimate that our approach is 5- to 10-fold cheaper than assembly-based methods, even excluding labor costs.

In spite of our success in identifying CREs that are active ubiquitously or in differentiated cell types, we have not yet demonstrated that this approach can efficiently identify the CREs of developmentally-regulated genes. The reasons for this failure could be trivial, such as an unfortunate selection of genes (or CREs) expressed at low levels. But there could also be substantive reasons, such as the very long range dispersal of the CREs of developmental genes, extending beyond the tens of kilobases that we scanned in this study, the presence of multiple shadow enhancers, each contributing a low level of activity in an additive manner, or the rapid evolution of these enhancers leading to low sequence conservation. Further studies could address this question by bringing in additional information, for example by looking at the long-range contacts of promoters with other parts of the genome using Hi-C (Lieberman-Aiden et al. 2009).

Despite these caveats, the resources we present here allow us to focus our search for CREs in the vast genome of *Parhyale* to a small subset of DNA fragments that are both accessible and functionally constrained, thus drastically reducing the search space. This approach has led us to discover robust ubiquitous, neuron- and muscle-specific regulatory elements, which will be valuable for building new genetic and imaging tools in this species. The same resources can now be used to identify CREs that are active in a range of other cell types in *Parhyale*. And the same approach should be applicable, with modest investment, to most organisms.

## Materials and Methods

### *Parhyale hawaiensis* culture and handling

We used the Chicago-F inbred line of *Parhyale hawaiensis* (Dana, 1853) in order to minimise polymorphisms and maximise mapping to the genome that has been sequenced from that line (Kao et al. 2016). The animals were cultured and embryos were collected in artificial sea water (specific gravity 1.022) using standard methods (Browne et al. 2005; Paris et al. 2022).

### Bulk ATAC-seq on whole embryos, embryonic and adult legs

The ATAC-seq protocols were based on (Buenrostro et al. 2013) (for embryos) and (Corces et al. 2017) (for embryonic and adult legs), with elements from the [10x Genomics Multiome protocol](https://cdn.10xgenomics.com/image/upload/v1745274143/support-documents/CG000375_DemonstratedProtocol_NucleiIsolationComplexSample_ATAC_GEX_Sequencing_Rev_D.pdf).

For whole embryo datasets E20, E23 and E24, single embryos at stages S20, S23 and S24 (Browne et al. 2005), respectively, were collected and placed in the lid of an eppendorf tube, in 50 µl ice-cold lysis buffer (10 mM Tris-HCl pH 7.4, 10 mM NaCl, 3 mM MgCl_2_, 0.1% Igepal CA-630). The egg membranes were removed, the cells were dissociated by gentle pipetting, and collected by centrifuging at 800 rcf for 10 min, at 4°C. The supernatant was discarded, the pellet was placed on ice and resuspended in 50 µl tagmentation mix, containing 25 μl 2x TD Buffer (Illumina #FC-121-1030) and 1.5 μl Tn5 Transposase (Illumina #FC-121-1030). Tagmentation was performed at 37°C for 30 minutes, shaking at 300 RPM.

For embryonic leg datasets EL1, EL3 and EL3, developing embryonic T4 to T8 legs were collected from pools of stage S25-26 embryos, when white eyes are just starting to be visible (Browne et al. 2005). The embryos were dissected in artificial sea water on a cold block; egg membranes were removed and 30-40 legs were collected in ice-cold buffer consisting of 10 mM Tris-HCl pH 7.4, 10 mM NaCl, 3mM MgCl_2_ and 1% bovine serum albumin (BSA). The legs were then placed in 500 µl ice-cold lysis buffer consisting of 10 mM Tris-HCl pH 7.4, 10 mM NaCl, 3mM MgCl_2_, 0.1% Igepal CA-630 (Merck #I3021), 1 mM DTT (Roche #10197777001), 0.01% Tween-20 (Sigma #P1379), and 0.001% digitonin (Promega #G944A), and the leg tissue was mechanically dissociated by 5 passes through a 25G x 5/8” needle using a 2.5 ml syringe. The lysate was incubated for 5 minutes on ice and then cuticle debris was removed by filtering with a 10 μm pluriStrainer cell strainer (pluriSelect #43-10010). The nuclei were collected by centrifuging at 4°C for 5 minutes at 500 rcf, washed in 400 µl ice-cold wash buffer (10 mM Tris-HCl pH 7.4, 10 mM NaCl, 3mM MgCl_2_, 1% BSA, 0.1% Tween-20, 1mM DTT), and collected again by centrifuging at 4°C for 5 minutes at 500 rcf. 18 to 25 thousand nuclei per sample, measured by staining a fraction of the suspension with trypan blue and counting the stained nuclei on a hemocytometer. Tagmentation was performed in 50 µl tagmentation mix, containing 25 µl 2X tagmentation buffer (Diagenode #C01019043) and 2.5 µl Tn5 loaded transposase (Diagenode #C01070012), at 37°C for 30 minutes, shaking at 300 RPM.

For adult leg datasets L1, L2, and L3, uninjured T4 and T5 legs were collected by anaesthetising adult animals in 0.02% clove oil in artificial seawater and cutting the legs at the proximal end of the merus (sample L1 and L2) or the basis (sample L3); 28 legs were collected for L1 and L3, 12 legs were collected for L2. Nuclei were isolated on a cold block, by opening each leg longitudinally and scraping the tissue from the cuticle using a surgical knife (Fine Science Tools #10316-14), followed by pipetting 25 times through a P200 micropipette tip in 250 µl of a lysis buffer consisting of 10 mM Tris-HCl pH 7.4, 10 mM NaCl, 3mM MgCl2, 1% BSA, 0.1% Igepal CA-630, 0.1% Tween-20, and 0.01% digitonin (for samples L1 and L3), or by 5 passes through a 25G x 5/8” needle using a 2.5 ml syringe (for sample L2). An equal volume of wash buffer (10 mM Tris-HCl pH 7.4; 10 mM NaCl; 3mM MgCl2; 1% BSA; 0.1% Tween-20) was then added and cuticle pieces were removed by passing the lysate through a 10 μm pluriStrainer cell strainer (pluriSelect #43-10010). Nuclei were collected by centrifuging at 4°C for 5 minutes at 500 rcf (samples L1 and L2) or 10 minutes at 600 rcf centrifugation (sample L3). 50 to 90 thousand nuclei were recovered per sample, measured on a hemocytometer as described earlier. Tagmentation was performed in 50 µl tagmentation mix, containing tagmentation buffer (Illumina #15027866 for sample L1 and L3, or Diagenode #C01019043 for sample L2) and 2.5 µl Tn5 loaded transposase (Illumina #15027865 for samples L1 and L3, or Diagenode #C01070012 for sample L2) at 37°C for 30 minutes, shaking at 300 RPM.

In parallel, we generated ATAC-seq data on genomic DNA from *Parhyale hawaiensis* that had been purified to remove histones and other proteins (’naked DNA’), to reveal intrinsic biases associated with tagmentation of DNA sequence rather than chromatin. The genomic DNA was purified as described for *P. aquilina* and *plumicornis* (below). Tagmentation was performed using the Nextera XT DNA Library kit (Illumina #FC-131-1024) according to the manufacturer’s recommendations and then sequenced on a NextSeq500 (PE 2×76).

For all samples, the tagmented DNA was purified using the Qiagen MinElute kit (Qiagen #28204) and amplified by a total of 9-17 PCR cycles using Nextera XT Index kit v2 primers (5 initial cycles followed by qPCR reaction to determine the total number of PCR cycles needed for each library). After PCR amplification, the libraries were purified by AMPure XP beads (Beckman Coulter) and were resuspended in 20 μl nuclease-free water. Sequencing was carried out on an Illumina NextSeq 500 sequencer; we carried out single-end 76 bp sequencing for the first sample we generated (E24), and then switched to paired-end 76 bp sequencing for the other samples, because this leads to more specific read mapping. The raw data (sequencing reads) of all the bulk ATAC-seq experiments are provided in Suppl. Data 1.

Reads were mapped to the *P. hawaiensis* genome (Phaw_5.0, GenBank GCA_001587735.2; updated in Almazán et al. 2022) using bowtie2 version 2.5.4 (Langmead and Salzberg 2012), using very-sensitive-local parameters. Mapped reads were filtered to remove mitochondrial reads and low quality mapped reads (MAPQ≥20), using SAMtools version 1.21 (Danecek et al. 2021) and BEDtools version 2.31.1 (Quinlan and Hall 2010), and duplicates using Picard MarkDuplicates version 2.27.5 (https://broadinstitute.github.io/picard/). ATAC-seq peaks were then identified using MACS2 version 2.2.7.1 (Zhang et al. 2008).

### Single-nucleus ATAC-seq on adult legs

Single-nucleus ATAC-seq was performed using the 10x Genomics single-cell capturing system. Libraries were prepared using the Chromium Next GEM Single Cell ATAC Reagent Kits v1.1 according to the manufacturer’s instructions (User Guide Rev. C). Nuclear suspensions from adult *Parhyale* legs were prepared as described above for sample L3. After lysis and filtering, nuclei were collected by centrifuging at 4°C for 10 minutes at 600 rcf. Then, the nuclear pellets were resuspended in 7 μl of 1X Nuclei Isolation Buffer provided by the kit. Nuclei suspensions containing 15,000 nuclei were used for the tagmentation reactions, which were then loaded on the 10x Genomics Chromium Controller. Libraries were sequenced on Illumina NextSeq500 (72 bp for R1 and R2, according to 10X Genomics User Guide recommendations). SN1 data were generated with two sequencing rounds (one NextSeq Mid Output and one NextSeq High Output), resulting in a total of 675 million reads. For SN2, data were generated with one NextSeq High Output sequencing round, obtaining a total of 404 million reads. Sequencing data were processed using the Cell Ranger ATAC software (v1.2.0, 10x Genomics, Pleasanton, CA) to perform alignment, barcode filtering, and peak calling following the standard pipeline. The raw data (sequencing reads) of the snATAC-seq experiments are provided in Suppl. Data 1. Reads were mapped to the same genome assembly as the bulk ATAC-seq data.

### Analysis of snATAC-seq data

Analyses were performed using the Signac v1.16.0 and Seurat v5.4.0 R packages (Stuart et al. 2021; Hao et al. 2024). Normalization and linear dimension reduction were performed using reciprocal latent semantic indexing (LSI) (Cusanovich et al. 2015), with the Seurat functions RunTFIDF, FindTopFeatures (with the min.cutoff parameter set to “q0”) and RunSVD, using default parameters unless otherwise specified. Datasets were integrated using reciprocal LSI projection (Stuart et al. 2019). Anchor sets were identified using the first 30 reduction axes, except the first one that strongly correlates with sequencing coverage. UMAP and cell clustering were done on the integrated dataset using (LSI) reduction (again using the 2:30 first axes of reduction). The two snATAC-seq datasets (SN1 and SN2) were processed in the same fashion.

Differentially accessible (DA) peaks were identified using the Seurat function FindAllMarkers with the parameter min.pct = 0.1. We noticed that all DA regions from cluster 19 were also open in cluster 7 and most DA regions from clusters 4, 5 and 7 were also open in cluster 19. We therefore suspect that cluster 19 may contain barcodes with doublets from clusters 4, 5 and/or 7; since it contains only 95 barcodes, we decided to remove it from the dataset in subsequent analyses.

Cell type labels on the snATAC-seq clusters were obtained by comparing snATAC-seq signal in gene bodies and promoter regions with gene expression in snRNA-seq (Almazán et al. 2022), using the Seurat function TransferData. Cell labels were transferred with good confidence (prediction score above 80%) for several cell types including neurons, muscles, or blood (Suppl. Figure 2).

### ATAC-seq data analysis and visualisation

Genome browser plots for bulk ATAC-seq, snATAC-seq and naked DNA were made with pyGenomeTracks (version 3.9; Lopez-Delisle et al. 2021) and normalised by counts per million. Cell-type-specific snATAC-seq tracks were made by combining the data of the two snATAC-seq experiments. The genome browser tracks of bulk and single nucleus ATAC-seq experiments are provided in Suppl. Data 2 and 3.

To assess the distribution of ATAC-seq peaks across genomic features (Figure 1B, Suppl. Table 2), genomic regions were classified as transcription start sites (TSS), exons, introns and intergenic regions. TSS was defined as the first base of annotated transcripts that contain at least one intron. Intronless transcripts were omitted to exclude poorly annotated transcripts from the analysis. Exonic regions were extracted from the same set of transcripts, excluding the TSS coordinate. Intronic regions were defined as the remaining regions of these transcripts that did not overlap with either the exons or TSS. Lastly, intergenic regions were defined as all remaining genomic regions, not assigned to TSS, exon, or intron. Unknown/unresolved nucleotides (Ns) in the genome and intron-less transcript regions were excluded from all feature sets.

ATAC-seq peak coordinates were determined from the MACS2 narrowPeak files for each dataset. A consolidated list of ATAC-seq peaks was generated by merging the peaks from all bulk ATAC-seq dataset using the Bedtools merge function version 2.31.1 (Quinlan and Hall 2010). Fragment (paired-read) counts per peak were quantified using featureCounts version 2.0.6 (Liao et al. 2014). Quantification was performed separately on the single-end sample (E24) and the paired-end sequenced samples (all other samples), and the resulting count matrices were combined. This consolidated peak set was used for assessing the overlap between ATAC-seq and cross-species islands of sequence conservation (Figure 2C-D).

The merged count matrix was normalised using variance stabilising transformation (VST) from DESeq2 version 1.48.2 (Love et al. 2014). Principal Component Analysis (Figure 1C) was performed on the VST-transformed counts using prcomp (stats version 4.5.2) with scaling enabled, and visualised using ggplot2 version 4.0.1 (Wickham 2016).

To generate the TSS enrichment heatmaps (Figure 1D), we normalised the ATAC-seq signal across datasets, the MACS2 bedgraph files representing the fragment coverage across the genome were scaled by fragments per million. For each dataset, the total number of fragments after mapping and filtering was divided by one million and the bedgraph signal values were divided by this scaling factor. Scaled bedgraph files were converted to bigwig format using bedGraphToBigWig version 377 (Kent et al. 2010). A signal matrix was generated using computeMatrix from deeptools version 3.5.6 (Ramírez et al. 2016) in reference-point mode, centred on TSS, with a bin size of 10 bp and +/- 1 kb flanking regions. Heatmaps were visualised using deeptools plotHeatmap, with the colour scale capped at the 98th percentile across all datasets. TSSs were sorted by mean signal intensity for each row in dataset EL3, which was the dataset with the highest FRiP score.

### Field collection and identification of *Parhyale* species

*Parhyale plumicornis* (Heller, 1866) were collected on the island of Milos, Greece, in rocky shores near Klima (coordinates 36.736594°N 24.417607°E, and N 36.732361°N 24.422056°E). Live adults were examined under a Nikon field microscope for preliminary identification based on the presence of the characteristic dense arrays of plumose setae on the ventral side of the second antennae, notably on articles 4 and 5 of the peduncle and on the first articles of the flagellum. Identification was then corroborated by additional characters scored on adults cultured in the laboratory, including a smooth body, length of second antenna about 2/3 of body length, inflated peduncle of the second antennae, subchelate first and the second gnathopods, robust and spinose pereopods, first uropod carrying a robust seta on the distal-lateral side of the peduncle, well defined inner ramus of the third uropod, which is not fused to the peduncle, and deeply cleft telson.

*Parhyale aquilina* (Costa, 1857) were collected in shallow water, in a well protected bay near Nea Peramos (Kavala), in Northern Greece (coordinates 40.825880°N 24.309377°E). Preliminary identification of adult males was based on the characteristic stout and inflated shape of the dactylus of the first pair of gnathopods, observed on a Nikon SMZ800N stereoscope. Females were selected based on the observation that they formed mating couples with those males. They could be identified more definitively as *P. aquilina* after genome sequencing (see below), based on sequence similarity with a previously sequenced gene fragment from this species (Iaciofano and Lo Brutto 2017).

*Parhyale darvishi* Momtazi & Maghsoudlou, 2016, were collected by hand between cobbles in the intertidal zone of Chabahar Bay (25.352533°N 60.598494°E), Gulf of Oman, placed in 99% ethanol and stored at -20°C. Preliminary identification of the adult males was conducted on a Nikon stereomicroscope based on the following diagnostic features: first antenna lacking dense setae, a stout second gnathopod, and an anterodistal tuft of setae on the maxillipedal palp. The identification was confirmed microscopically by comparing diagnostic characters with the original species description (Momtazi and Maghsoudlou 2016), including the absence of setae on the posterior margin of the propodus of pereopods 6 and 7, the presence of five apical robust setae on the outer ramus of uropod 3, and two robust setae on the palm of the first male gnathopod, with the proximal seta being distinctly incurved. Female specimens were selected based on their association with males in the field and confirmed in the laboratory by examining their diagnostic features under a microscope: a transverse palm defined by two stout setae, a medial process on the posterior margin of the palm of the second gnathopod, and oostegites bearing curl-tipped setae.

### Low coverage genome sequencing and cross-species read mapping

Genomic DNA was extracted from single individuals that were flash frozen in liquid nitrogen, using the NucleoSpin Tissue kit (Macherey Nagel #740952) according to the manufacturer’s recommendations. Whole genome libraries were built for each species using the Nextera XT DNA Library kit (Illumina #FC-131-1024) according to the manufacturer’s recommendations and then sequenced on a NextSeq500 (PE 2×76). The raw data (sequencing reads) obtained from *P. aquilina*, *P. darvishi* and *P. plumicornis* are provided in Suppl. Data 4.

We noticed that one sequencing round on *P. plumicornis* included reads corresponding to a plasmid that was used in our laboratory, which includes 3 fragments from the genome of *P. hawaiensis* (the *PhOpsin1* promoter and two fragments from the *Dll-e* locus), representing a contamination of our library. We removed the contaminating reads by mapping the sequenced reads onto the plasmid sequence, using bowtie with the parameters “--end-to-end --very-sensitive”. We discarded the reads with matches to the plasmid sequence, except ones that had 2 or more mismatches to the plasmid (found in ≥2 reads). Overall a total of 37,385 reads were discarded from the sequencing output file, leaving 424,305,645 reads.

To estimate genome coverage, the reads sequenced from *P. aquilina*, *P. darvishi* and *P. plumicornis* were used to generate a genome assembly for each species using ABySS-pe v4.2.1 (Jackman et al. 2017), with parameters kc=2 k=25 n=2. The resulting assemblies were highly fragmented (N50 < 1kb), as expected considering the low sequencing coverage. Sequences corresponding to the crustacean BUSCO gene set (version odb12) were identified in these assemblies using BUSCO v6.0.0 (parameter --metaeuk) (Tegenfeldt et al. 2025). To estimate sequencing coverage for each species, we mapped the reads back to the genome assembly using bowtie2 v2.3.5.1 (Langmead and Salzberg 2012) with stringent parameters (--end-to-end --very-sensitive --score-min L,-0.2,-0.2). We discarded reads that harbored over two mismatches or a mapping quality below 30, and duplicate reads. We then computed the per base coverage on the BUSCO genes (distributions shown in Suppl. Figure 3). Estimates of genome size were obtained by dividing the number of sequenced bases by the mean coverage. The estimated coverage was 16x, 11x and 10x, corresponding to estimated genome sizes of 1.1 Gbp, 2.8 Gbp and 3.0 Gbp for *P. aquilina*, *P. darvishi* and *P. plumicornis*, respectively. As a control, we followed the same procedure for *P. hawaiensi*s, for which genome size is known (Parchem et al. 2010; Kao et al. 2016), and found a genome size of 3.0 Gbp instead of 3.6 Gbp (with a genome coverage of 5.8x). We cannot determine whether our predicted genome size is more accurate than the published estimate, or whether our method slightly overestimates coverage (and thus underestimates genome size). In any case, these results suggest that the genome sizes of *P. darvishi* and *P. plumicornis* are similar to that of *P. hawaiensis*, but the genome of *P. aquilina* is about 3 times smaller.

Reads from all species were mapped to a genome assembly of *P. hawaiensis* (Phaw_5.0, GenBank GCA_001587735.2; updated in Almazán et al. 2022), using bowtie2 v2.3.5.1 with the parameters “--very-sensitive-local --local -3 10”, and low quality mapping were removed using the function “view” from samtools v1.21 with the parameter ‘-q 10”. Conservation tracks were produced using the function pileup form macs2 with the parameter “--extsize 75”.

Genome browser plots for sequence conservation were generated with pyGenomeTracks (version 3.9; Lopez-Delisle et al. 2021) and normalised by counts per million, as previously described for ATAC-seq. The genome browser tracks showing the cross-species sequence conservation are provided in Suppl. Data 5.

Islands of sequence conservation were determined for each species as non-exonic genomic regions (TSS, introns and intergenic regions) with at least one mapped read; their lengths varied from 23 to 5771 nucleotides, with a median of 59 to 69 nucleotides). We considered that conserved sequence islands overlap between species if their coordinates on the *P. hawaiensis* genome are shared over at least 50% of the length of the shortest island (Figure 2B). We followed the same logic for assessing the overlap of ATAC-seq peaks with conserved regions (Figures 2C-D). For the latter analysis we used the consolidated list of ATAC-seq peaks (see above).

### Phylogeny and estimate of divergence times

The molecular phylogeny of the sequenced *Parhyale* species (Figure 3A) was generated based on the sequences of conserved single-copy BUSCO genes. We ran BUSCO with the same parameters as described earlier on three additional species with available genome assemblies: *P. hawaiensis* (Phaw_5.0, GenBank GCA_001587735.2; updated in Almazán et al. 2022), *Hyalella azteca* (NCBI RefSeq assembly GCF_000764305.2, Poynton et al. 2018) and *Daphnia pulex* (NCBI RefSeq assembly GCF_021134715.1). We selected the 23 BUSCO genes that were found in all 6 species, aligned each gene set separately using macse v2.07 that preserves coding frames (Ranwez et al. 2018), and concatenated the gene alignments using AMAS (Borowiec 2016). We selected the 29,097 sites that were represented in all four *Parhyale* species and in *H. azteca*. The concatenated alignment is available in Suppl. Data 6. The phylogeny was generated by the Maximum Likelihood approach using iqtree v3.0.1 (Wong et al. 2025) with the parameters -m MFP -bb 1000 -nt AUTO. Divergence times were estimated using iqtree v2.4.0 on the output tree and the same input alignment, calibrated against an estimated divergence of 110 to 140 million years between *Parhyale hawaiensis* and *Hyalella azteca* (Cannizzaro and Berg 2022),

### Identification of CREs using transgenic reporters

Our initial efforts to identify neuron-specific CREs (prior to availability of ATAC-seq and sequence conservation data) focused on candidate genes expressed in neurons or involved in neurogenesis. The reporter constructs included 1.5 to 5.5 kb of sequence immediately upstream of the start codon of each gene: reporters with 1.5 and 3.6 kb upstream sequence for *bruchpilot* (mikado.phaw_50.283862bG132); 2.3 and 4 kb for *VAChT* (MSTRG.41555); 2 kb for *synaptotagmin 4* (mikado.phaw_50.283817fG572); 4.2 kb for *achaete/scute* (MSTRG.45479); 4.8 kb for *senseless* (MSTRG.40789); 3.9 kb for *cut* (MSTRG.41143); 5.5 kb for *prospero* (MSTRG.45422); 5 kb for *neuralised* (MSTRG.5439). The sequences of these fragments are available in Suppl. Data 7. Based on our transcriptome assemblies (Kao et al. 2016; Sinigaglia et al. 2022) these fragments include the 5’UTR, the 5’ end of the annotated transcript (putative TSS) and 1 to 5 kb of sequences further upstream. These putative CREs were amplified by PCR on *Parhyale* genomic DNA and cloned immediately upstream of the coding sequence of EGFP and the SV40 early transcriptional terminator, and placed in a *Minos* transposon vector (Pavlopoulos and Averof 2005) that also carries *PhOpsin1-EGFP* as a transgenesis marker (Ramos et al. 2019). These reporter constructs were co-injected with *Minos* transposase mRNA into early *Parhyale* embryos, as described previously (Pavlopoulos and Averof 2005; Kontarakis and Pavlopoulos 2014); typically, the injection mix contained 250 ng/µl donor plasmid and 200 ng/µl Minos transposase mRNA. The injected embryos were raised in filtered artificial sea water containing 100 units/mL penicillin, 100 μg/mL streptomycin and 0.25 μg/mL amphotericin B (Gibco #15240096). The surviving embryos were screened for fluorescence on a Leica MZ16F stereoscope during mid and late embryonic stages. At least 50 embryos expressing the PhOpsin1-EGFP marker were scored per construct, but no additional expression was detected in the nervous system.

Candidate ubiquitous CREs were identified by combining information from snRNA-seq, snATAC-seq and sequence conservation. First, using the integrated snATAC-seq dataset, we selected the peaks that are accessible in at least 20% of the cells within every cell cluster and do not show differential accessibility across clusters (Seurat function FindAllMarkers p_val_adj > 0.5). Second, among those peaks, we selected the ones that are nearest a gene (using the Signac function ClosestFeature) that is expressed in at least 20% of the cells within each cell cluster, and is not differentially expressed across clusters (Seurat function FindAllMarkers p_val_adj >0.5, or pct.1>0.5 and pct.2>0.5 and abs(pct.1-pct.2) <0.1), based on the snRNA-seq data generated by (Almazán et al. 2022). Third, among the resulting peaks, we selected those that included regions of cross-species sequence conservation. Among the resulting peaks, we focused on the orthologues of *Drosophila* genes *headcase/hdc* (MSTRG.41953) and *muscleblind/mbl* (MSTRG.26075) (Figure 4A,B). To the *mbl* promoter region, we added an ATAC-seq peak with conserved sequences that is located within an intron >50 kb downstream of the promoter (Figure 4B). These candidate CREs were placed upstream of the coding sequence of the mNeonGreen fluorescent protein fused with histone H2B to drive nuclear localisation (Wolff et al. 2018), followed by the P10 transcriptional terminator (Pfeiffer et al. 2012), designated as *H2B-mNeonGreen-P10*, by gene synthesis (Twist Bioscience, USA), in the context of a *Minos* vector carrying the *PhOpsin1-EGFP* transgenesis marker (described above). The *mbl* promoter fragment includes a splice donor sequence and the 5’ end of the gene’s first intron, which carries ATG triplets that could be misread as start codons. To ensure that this region is spliced out, we added the 3’ end of an intron and splice acceptor from the *Parhyale Distal* locus (Kontarakis et al. 2011), prior to the mNeonGreen coding sequence. The annotated sequences of both CREs are available in Suppl. Data 7. These reporter constructs were injected into *Parhyale* embryos as described above, and screened in mid and late embryonic stages and in juveniles using a Zeiss Axiozoom V16 stereoscope. The image shown in Figure 4B’ was captured on a Zeiss LSM 800 confocal microscope equipped with a Plan-Apochromat 20 x/0.8 M27 objective (Zeiss 420650-9901-000).

Candidate neuron-specific CREs were identified by combining information from RNA-seq, ATAC-seq and sequence conservation. First, we selected the peaks in the integrated snATAC-seq dataset that are enriched in the cells predicted to be neurons compared to the cells predicted to be non-neuronal (with a confidence score ≥0.8, based on the label-transfer approach described earlier), using the Seurat function FindAllMarkers (test.use = ‘LR’,min.pct = 0.2) with the following filtering parameters: corresponding ATAC-seq reads should be found in at least 20% of the predicted neuronal cells, and in at least 10% more cells than in the other cell types combined, the log2 fold change enrichment should be over 3 and the adjusted p.value below 10^(-100). We only kept the peaks that were nearest to a gene labeled as neuronal-specific, using our snRNA-seq data (Almazán et al. 2022) and the ClosestFeature function from Signac and FindMarkers from Seurat. A list of 37 peaks near 33 genes was manually inspected on the IGV genome browser using our bulk and snATAC-seq and sequence conservation data, and reduced to a set of 7 genes, including 7 promoter regions and 5 additional associated CREs. Promoter regions included the sequence from the start of the ATAC-seq peak up to the nucleotide before the start codon; associated CREs included the sequence of the entire ATAC-seq peak. Reporter constructs were generated by placing the candidate CRE and promoter sequences upstream of the coding sequences of beta galactosidase and EGFP, separated by the T2A ribosome skipping peptide, and the SV40 early transcriptional terminator. These reporters were placed in a *Minos* vector carrying the *PhOpsin1-EGFP* transgenesis marker (as described above). The reporter constructs were generated by gene synthesis (Twist Bioscience, USA) and conventional subcloning. The annotated sequences are available in the Suppl. Data 7. The activity of these reporters was tested by microinjecting *Parhyale* embryos as described above. Approximately 400-700 embryos were injected per construct. The surviving embryos were screened for fluorescence on Leica MZ16F and Zeiss AxioZoom V16 fluorescence stereoscopes during stages S20 to S28. For each construct, we found that 20-30% of injected embryos surviving to late embryonic stages expressed the *PhOpsin1-EGFP* transgenesis marker, either unilaterally or bilaterally (10 to 80 positive embryos per construct). In addition to this marker, approximately 20% of the embryos injected with the *neuro5* and *neuro6* constructs showed strong fluorescence in the brain and ventral nerve cord.

A second set of constructs were made in which the *neuro5* and *neuro6* regulatory elements are used to express membrane-localised fluorescent proteins (using the Src64B and HRas tags; Karapidaki et al. 2024), to label neuronal projections. This includes two constructs carrying *neuro5-Src64B-mNeonGreen* and *neuro6-Src64B-mNeonGreen* (followed by the P10 transcriptional terminator; Pfeiffer et al. 2012), and a construct carrying both *neuro6-mScarlet3-HRas* (followed by the SV40 early transcriptional terminator) and *neuro5-Src64B-mNeonGreen* (followed by the P10 transcriptional terminator). The first two were cloned in a *Minos* vector carrying the *PhOpsin1-EGFP* transgenesis marker (see above); the latter was cloned in a *Minos* vector carrying no additional marker. These constructs were microinjected and screened as described above. Mosaic G0 embryos were imaged on a Zeiss LSM 800 confocal microscope equipped with a Plan-Apochromat 20 x/0.8 M27 objective (Zeiss 420650-9901-000).

To identify muscle-specific transgenic line, we focused on the *Myosin heavy chain (Mhc)* gene of *Parhyale* (MSTRG.35247), which is specifically and strongly expressed in all muscle cells in the snRNA-seq data of (Almazán et al. 2022). We identified the promoter of *Mhc* and two putative CREs that are accessible specifically in muscle nuclei and contain regions that are conserved across *Parhyale* species. The sequences of these putative CREs are available in Suppl. Data 7. These sequences were placed upstream of the coding sequence of mNeonGreen followed by the P10 transcriptional terminator, by gene synthesis (Twist Bioscience, USA), in the context of a *Minos* vector carrying the *PhOpsin1-EGFP* transgenesis marker (as described above). The activity of these reporters was tested by microinjection into *Parhyale* embryos, as described above. Late embryos (from S26) were screened for both *Mhc* and *PhOpsin1* activity using a Leica MZ16F fluorescence stereoscope.

To identify CREs regulating the activity of leg patterning genes, we explored ATAC-seq peaks with sequence conservation around the genes *dll-e*, *dac1* and *dac2,* corresponding to genes MSTRG.27394, MSTRG.33907 and MSTRG.33908 in the genome annotation of (Sinigaglia et al. 2022). Putative CREs and promoter regions, identified by the TSS peak, were placed upstream of *H2B-mNeonGreen-P10*, within the *Minos* transposon vector containing the *PhOpsin1-EGFP* transgenesis marker, as described earlier. The CREs were tested by microinjecting *Parhyale* embryos, as described above. Embryos were screened for CRE activity every day from the start of leg development (S19) to hatching, and *PhOpsin1* activity was scored at the final developmental stages. CREs were considered inactive, or below our detection threshold, when at least 10 *PhOpsin1* positive embryos were identified while embryos did not consistently show the expected distal or mid leg expression pattern. Long and short versions of the promoter region were tested for *Dlle* and *dac2*. Not all obvious ATAC-seq peaks around *Dlle* were tested since both long and short versions of the promoter alone gave weak ubiquitous expression. The sequences of these putative CREs are available in Suppl. Data 7.

## Data availability

The sequencing data of the bulk and single-nuclei ATAC-seq experiments are available at NCBI GEO accessions GSE325175 and GSE325176, respectively. The NGS sequencing data from the genomes of *P. aquilina*, *P. darvishi* and *P. plumicornis* are available at NCBI BioSample accessions SAMN56533921, SAMN56533922 and SAMN56533923, respectively. The supplementary data files are available at https://zenodo.org/records/19020963.

## Acknowledgements

We thank Oliver Coleman of the Natural History Museum in Berlin for advice on collecting *Parhyale*, Pavlos Vidoris for support during field work, Chiara Sinigaglia for advice on leg staging and dissection, Alba Almazan for advice on nuclear isolation, and Benjamin Gillet and Sandrine Hughes of the IGFL sequencing platform for support with Illumina sequencing. This research was supported by grants from the Fondation pour la Recherche Médicale (grant EQU202303016278) and the European Research Council (grant ERC-2015-AdG 694918), a doctoral fellowship from Boehringer Ingelheim Fonds, and a fellowship from the Marie Curie ITN programme EvoCell (H2020-MSCA-ITN-2017 #766053), under the European Union Horizon 2020 programme. SLB was supported by the National Recovery and Resilience Plan of the European Union (project CN_00000033, CUP B73C22000790001).

## Author contributions

MA and MP conceived and supervised the project; GF, ES and ÇÇ generated the bulk ATAC-seq data; GF analysed the bulk ATAC-seq data; ES generated and analysed the single-nucleus ATAC-seq data; MP sequenced the genomes, determined the genome-wide patterns of sequence conservation, performed the phylogenetic analysis and supervised the bioinformatic analyses; IK and GF optimised the transgenic reporter approach; IK, SM and MD identified the neuron-specific, muscle-specific and ubiquitous CREs, respectively; GF tested the CREs of developmental genes; FM collected and identified *P. darvishi*; MA and CA collected *P. aquilina*; SLB provided advice for field collections and identified *P. plumicornis*; MA collected *P. plumicornis*; MA drafted the first version of the manuscript; all the authors edited the manuscript and contributed to the figures.

## Supplementary Materials

## Supplementary Tables

**Supp. Table 1.**
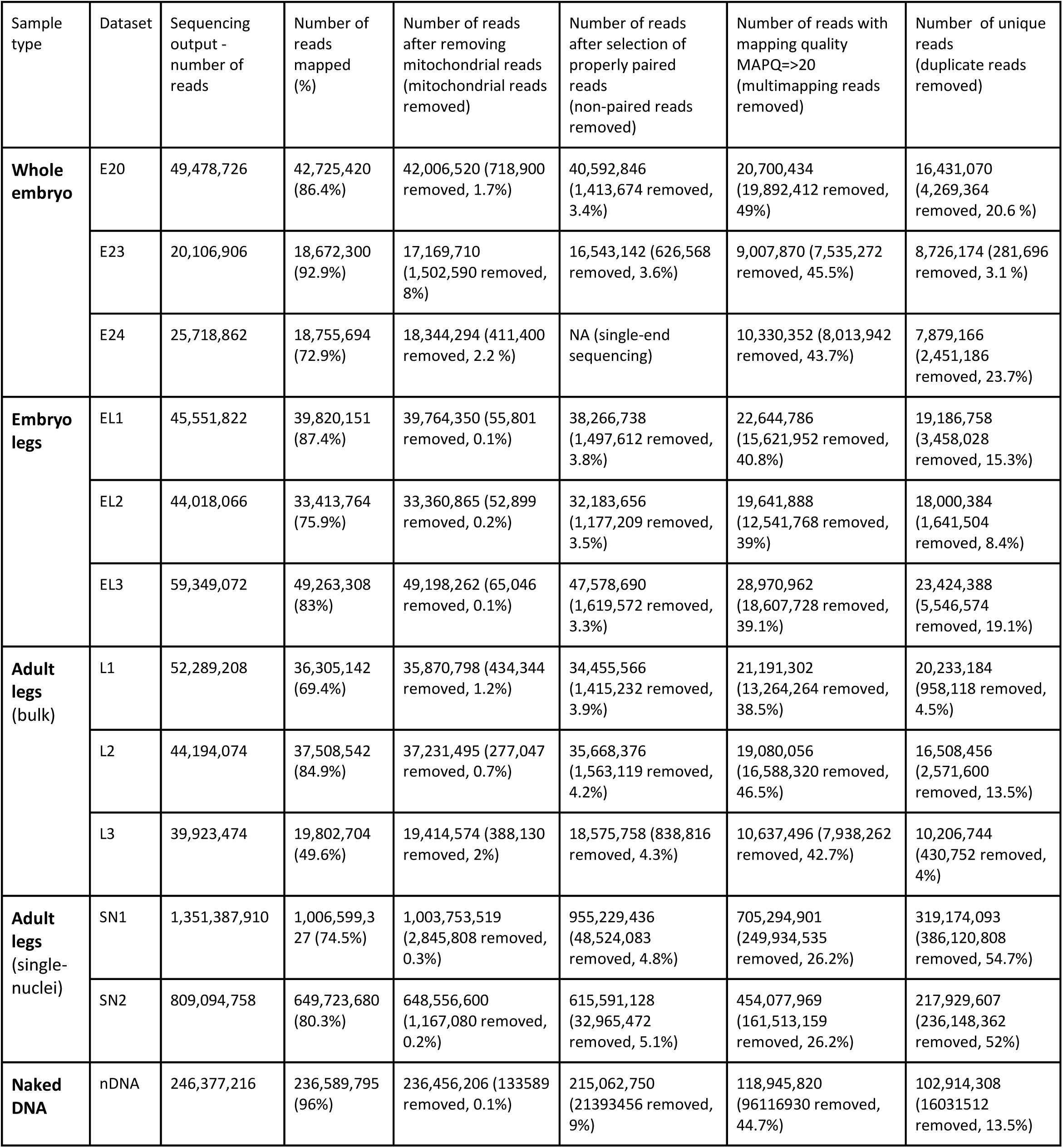
Read mapping of ATAC-seq data to the *Parhyale hawaiensis* genome. Statistics of ATAC-seq mapping using Bowtie2 to the *P. hawaiensis* genome, and subsequent filtering to remove mitochondrial reads, unpaired reads (except E24, which was sequenced single end), low mapping quality reads, and PCR duplicated reads. Paired-end reads were counted as two reads.

**Suppl. Table 2.** Number of nucleotides in ATAC-seq peaks in TSS, exon, intron and intergenic sequences. We determined the nucleotides of ATAC-seq peaks belonging to each of these genomic features: transcription start sites (TSS), exons, introns, and intergenic regions. Counts on TSS and exons were also determined for intron-containing genes only. Undetermined nucleotides (Ns) in the genome assembly were excluded from the counts.

| Sample type | Dataset | All TSS<br>(56,307 nucl) | TSS in intron-containing genes<br>(27,955 nucl) | All Exons<br>(129,980,737 nucl) | Exons in intron-containing genes<br>(94,685,505 nucl) | Introns<br>(673,377,575 nucl) | Intergenic<br>(1,390,661,853 nucl) |
| --- | --- | --- | --- | --- | --- | --- | --- |
| Whole embryo | E20 | 2,820 | 2,142 | 3,434,318 | 1,551,895 | 4,184,806 | 10,599,571 |
|  | E23 | 3,158 | 2,669 | 2,368,108 | 1,256,933 | 3,680,699 | 10,594,791 |
|  | E24 | 5,842 | 5,001 | 4,480,014 | 3,142,237 | 10,417,667 | 22,338,694 |
| Embryo legs | EL1 | 5,163 | 4,260 | 4,608,493 | 2,682,977 | 9,419,742 | 15,523,581 |
|  | EL2 | 5,161 | 4,421 | 3,808,532 | 2,242,178 | 8,236,839 | 13,758,216 |
|  | EL3 | 5,832 | 4,865 | 5,816,624 | 3,698,369 | 12,442,011 | 20,580,140 |
| Adult legs | L1 | 6,193 | 5,360 | 4,212,182 | 2,470,145 | 9,391,666 | 15,678,750 |
|  | L2 | 4,066 | 3,301 | 3,265,858 | 1,521,817 | 4,817,687 | 9,726,381 |
|  | L3 | 4,343 | 3,821 | 2,558,498 | 1,378,514 | 4,231,107 | 8,936,821 |
| Naked DNA | nDNA | 5,551 | 2,496 | 12,311,372 | 6,304,430 | 33,198,617 | 70,905,248 |

**Suppl. Table 3.**
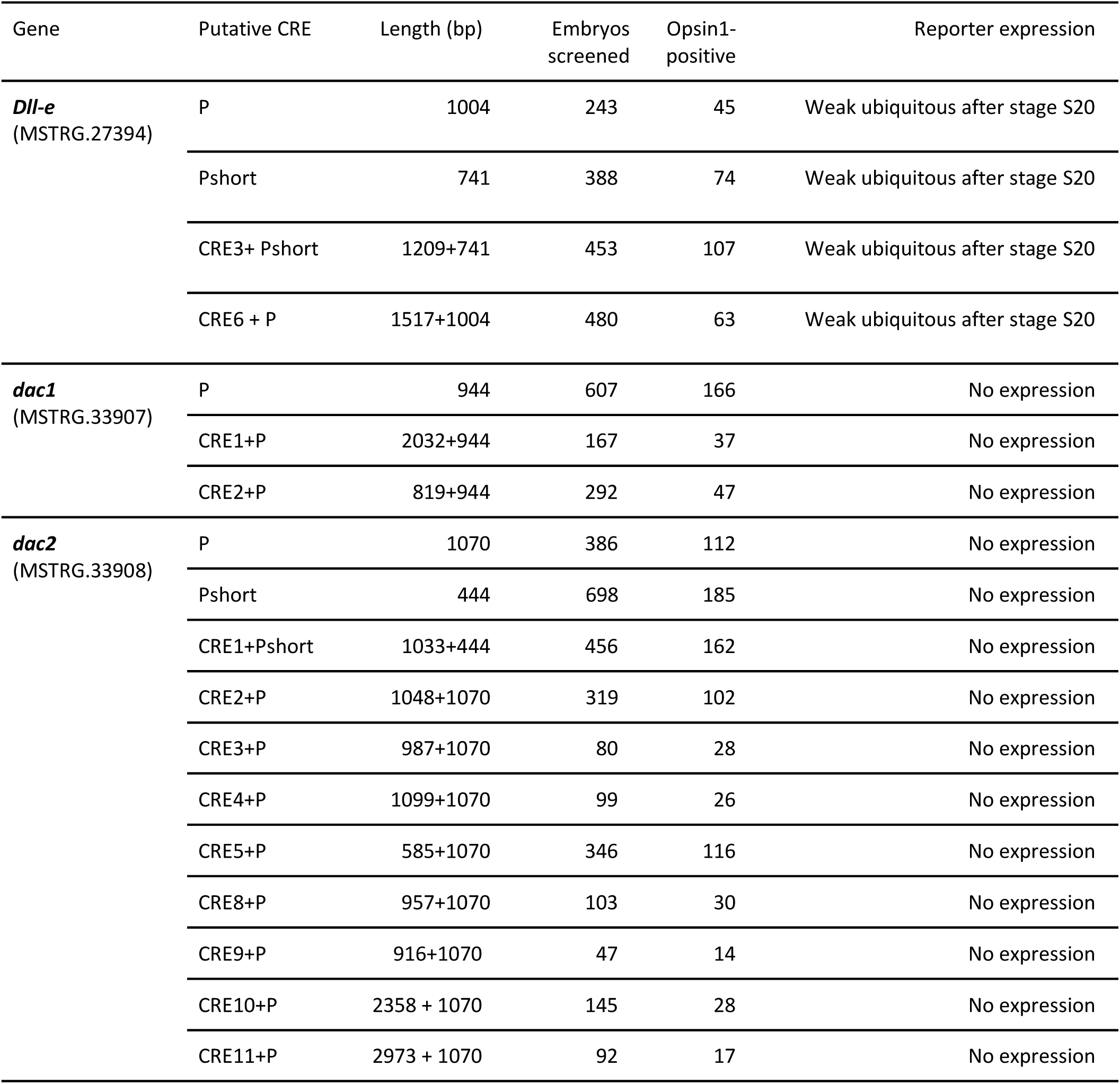
Putative CREs of *Parhyale* developmental genes tested using transgenic reporters. The transgenic reporters tested carried the promoter region (P) of the gene of interest and additional putative CREs upstream of the mNeonGreen coding sequence. The fragments tested are shown in Suppl. Figure 6. In 6-10% of the embryos screened for *Dll-e* activity we observed expression patterns that were not reproducible; we interpret these as enhancer traps associated with specific sites of transgene insertion, rather than the activity of sequences carried in the reporter. The P and CRE sequences are given in Suppl. Data 7.

| Gene | Putative CRE | Length (bp) | Embryos screened | Opsin1-positive | Reporter expression |
| --- | --- | --- | --- | --- | --- |
| ***Dll-e***<br>(MSTRG.27394) | P | 1004 | 243 | 45 | Weak ubiquitous after stage S20 |
|  | Pshort | 741 | 388 | 74 | Weak ubiquitous after stage S20 |
|  | CRE3+ Pshort | 1209+741 | 453 | 107 | Weak ubiquitous after stage S20 |
|  | CRE6 + P | 1517+1004 | 480 | 63 | Weak ubiquitous after stage S20 |
| ***dac1***<br>(MSTRG.33907) | P | 944 | 607 | 166 | No expression |
|  | CRE1+P | 2032+944 | 167 | 37 | No expression |
|  | CRE2+P | 819+944 | 292 | 47 | No expression |
| ***dac2***<br>(MSTRG.33908) | P | 1070 | 386 | 112 | No expression |
|  | Pshort | 444 | 698 | 185 | No expression |
|  | CRE1+Pshort | 1033+444 | 456 | 162 | No expression |
|  | CRE2+P | 1048+1070 | 319 | 102 | No expression |
|  | CRE3+P | 987+1070 | 80 | 28 | No expression |
|  | CRE4+P | 1099+1070 | 99 | 26 | No expression |
|  | CRE5+P | 585+1070 | 346 | 116 | No expression |
|  | CRE8+P | 957+1070 | 103 | 30 | No expression |
|  | CRE9+P | 916+1070 | 47 | 14 | No expression |
|  | CRE10+P | 2358 + 1070 | 145 | 28 | No expression |
|  | CRE11+P | 2973 + 1070 | 92 | 17 | No expression |

## Supplementary Figures

**Suppl. Figure 1.**
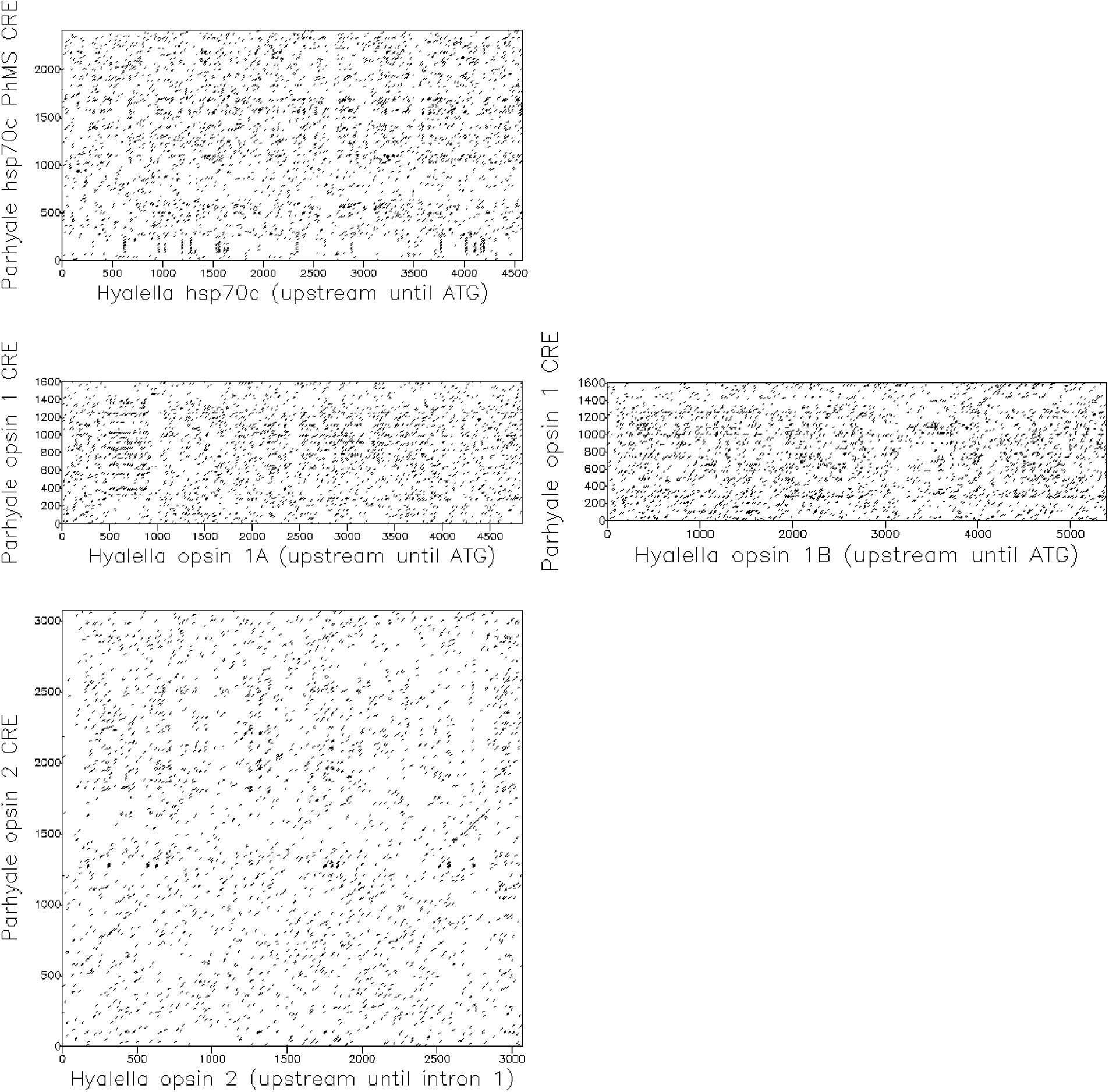
Sequence comparisons of known *Parhyale* CREs with homologous regions from Hyalella azteca. Dot plot alignments between previously characterised *Parhyale* regulatory fragments from the hsc70 (PhMS CRE, Pavlopoulos and Averof 2005), opsin 1 and opsin 2 genes (Ramos et al. 2019) and homologous regions of the *Hyalella azteca* genome (sequence accessions NW_025942174, NW_025945614 and NW_025930472, respectively; Poynton et al. 2018). Both putative opsin 1 genes of *Hyallella* were tested. The dot plots were made using the EMBOSS Dotmatcher tool (Rice et al. 2000), with low stringency settings (window size 10, threshold 30). Regions of sequence conservation should appear as diagonal lines. The only region of significant sequence similarity, in opsin 2, coincides with part of the coding sequence.

**Suppl. Figure 2.**
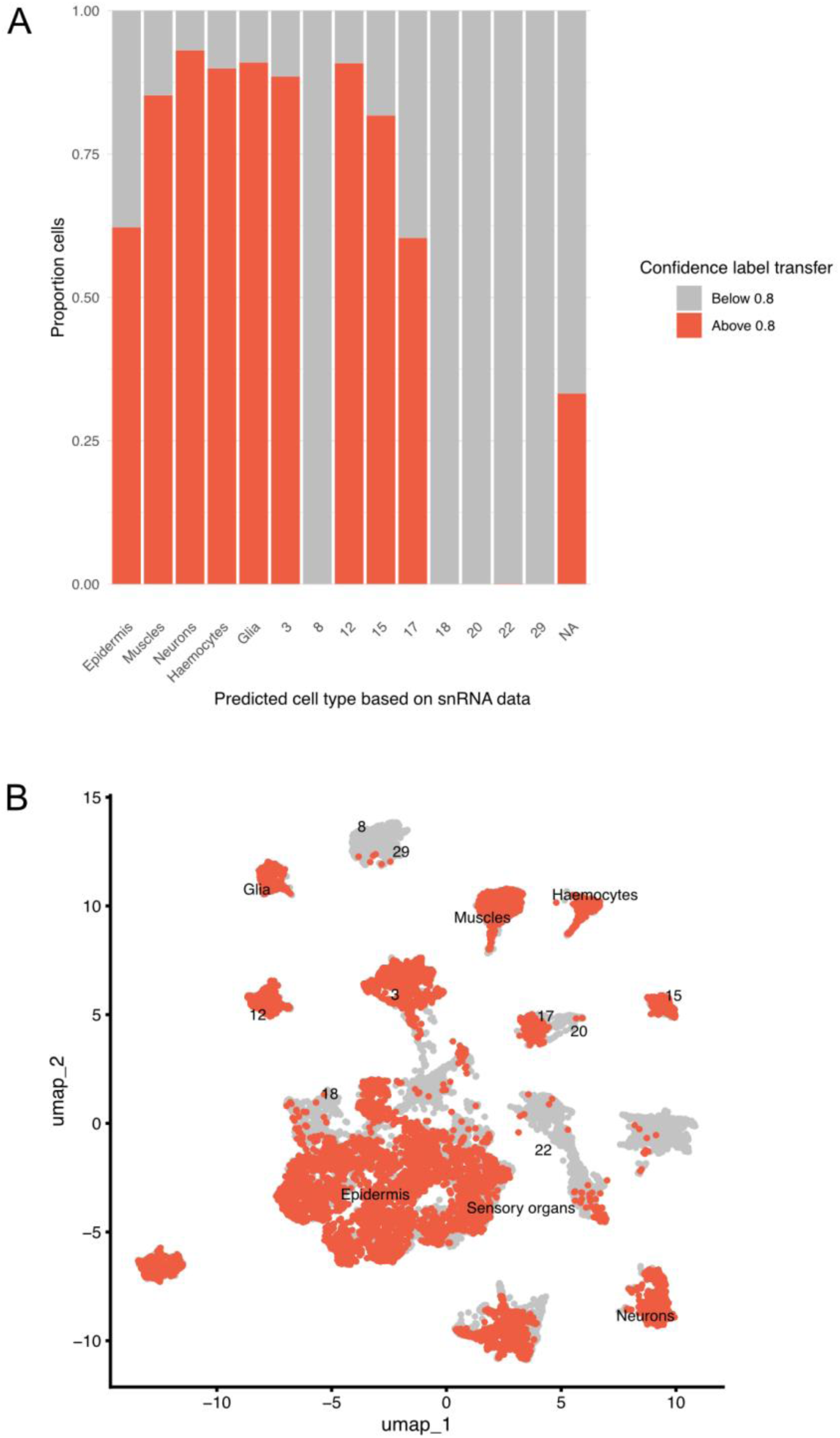
Label transfer scores from snRNA-seq to snATAC-seq. (A) Distribution of scores for the transfer of cell labels from the snRNA-seq (Almazán et al. 2022) to snATAC-seq (SN1 and SN2) data. (B) UMAP of the snATAC-seq experiments, color-coded according to the label transfer scores. Both panels show that cell identities of several cell types (including muscles, neurons, epidermis and haemocytes) can be transferred from RNA-seq to ATAC-seq clusters with relatively high confidence from the RNA-seq data.

**Suppl. Figure 3.**
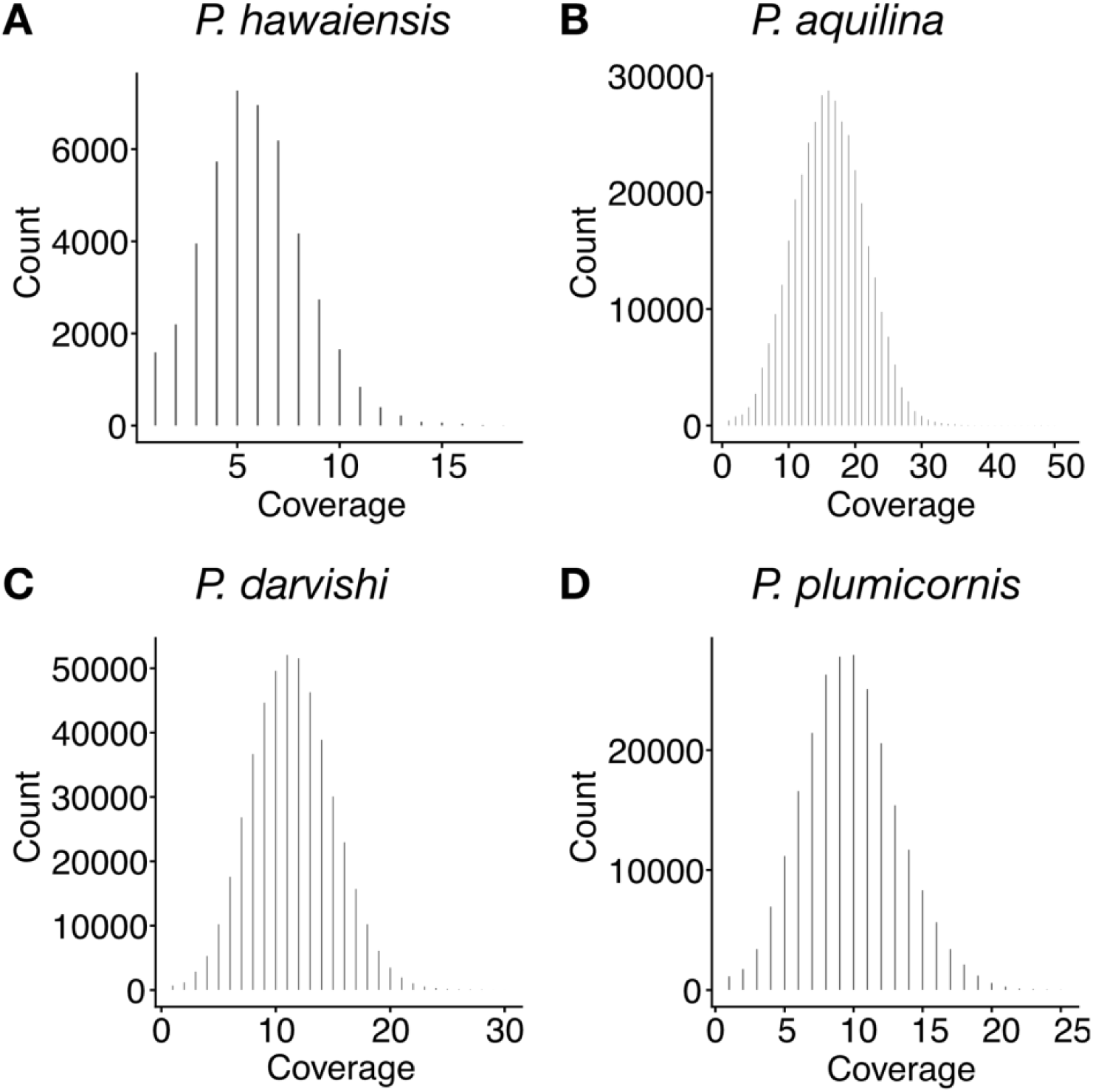
Estimates of genome coverage in *P. aquilina*, *P. darvishi* and *P. plumicornis*. Distribution of per nucleotide sequence coverage, based on the sequences of single-copy BUSCO genes in *P. hawaiensis*, *P. aquilina*, *P. darvishi* and *P. plumicornis* (see Methods).

**Suppl. Figure 4.**
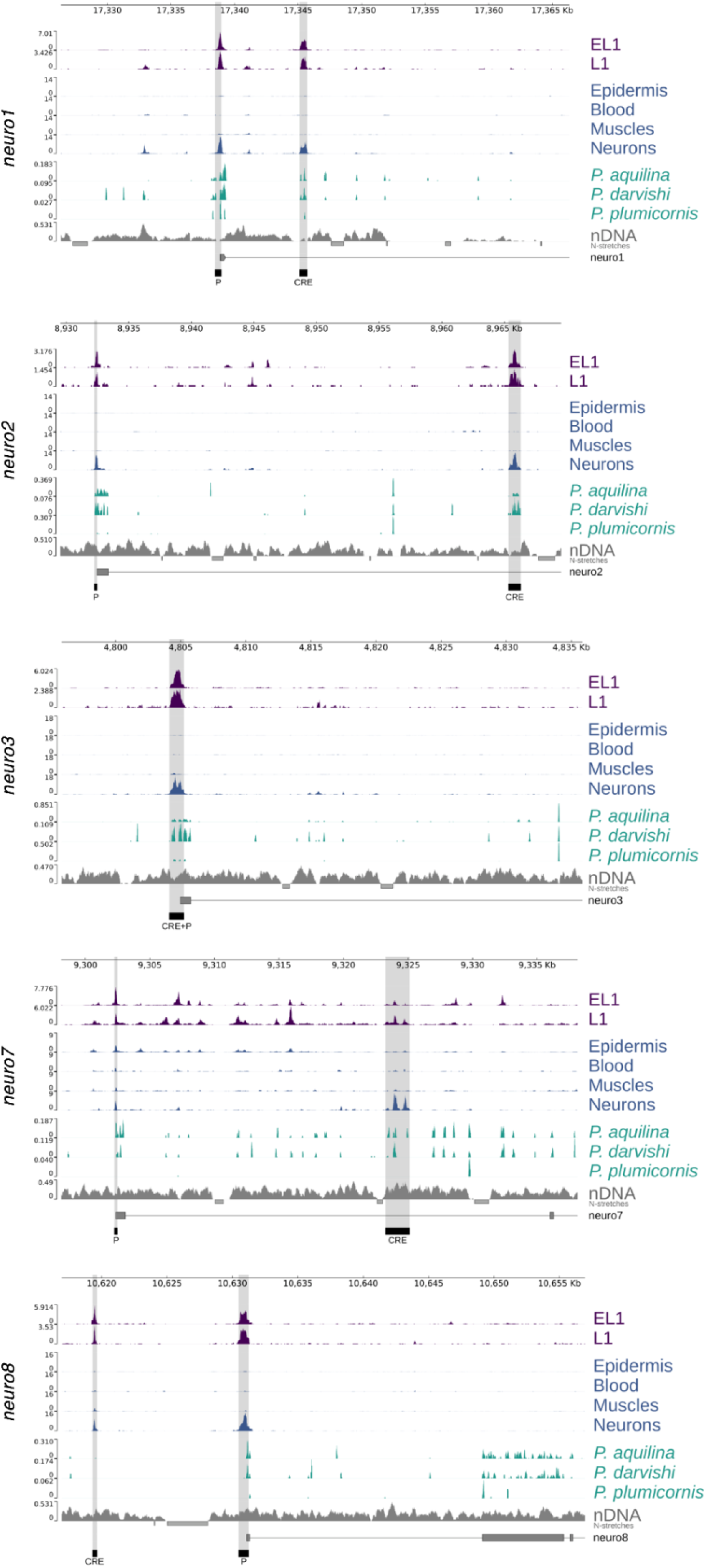
Putative CREs of *Parhyale* neuron-specific genes tested using transgenic reporters. Genome browser plots for 5 loci harboring putative neuron-specific CREs (highlighted in grey). The tracks of the other two loci that we tested, *neuro5* and *neuro6*, are shown in Figure 5A,B.

**Suppl. Figure 5.**
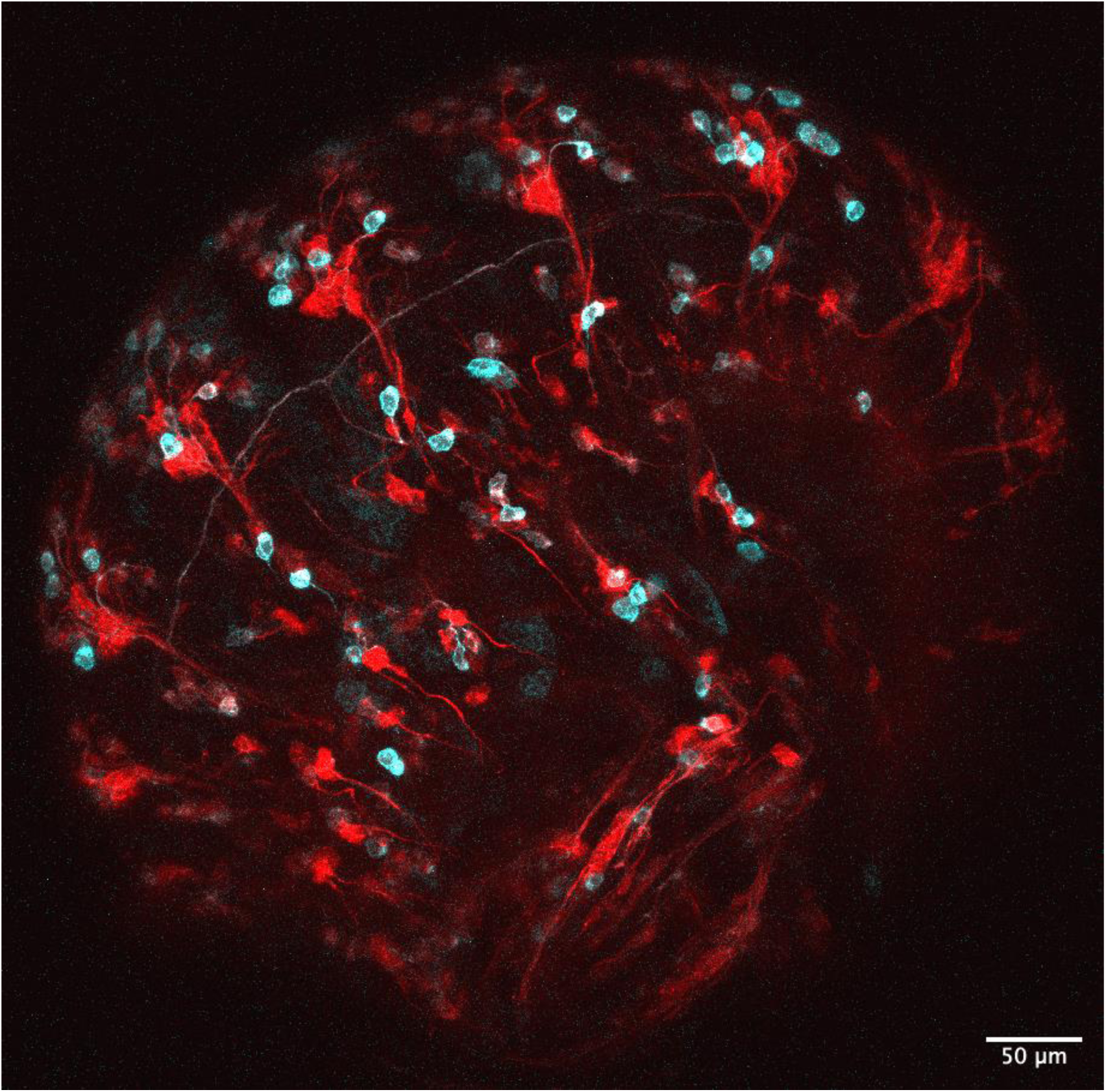
Comparison of the activity of *neuro5* and *neuro6* reporters. Dorso-lateral view of a transgenic embryo carrying the *neuro5>Src64B-mNeonGreen* (in cyan) and *neuro6-mScarlet3-HRas* (in red) transgenes. Dorsal side is located towards the top-left. Max projection of image stack captured by confocal microscopy. Scale bar, 50 µm.

**Suppl. Figure 6.**
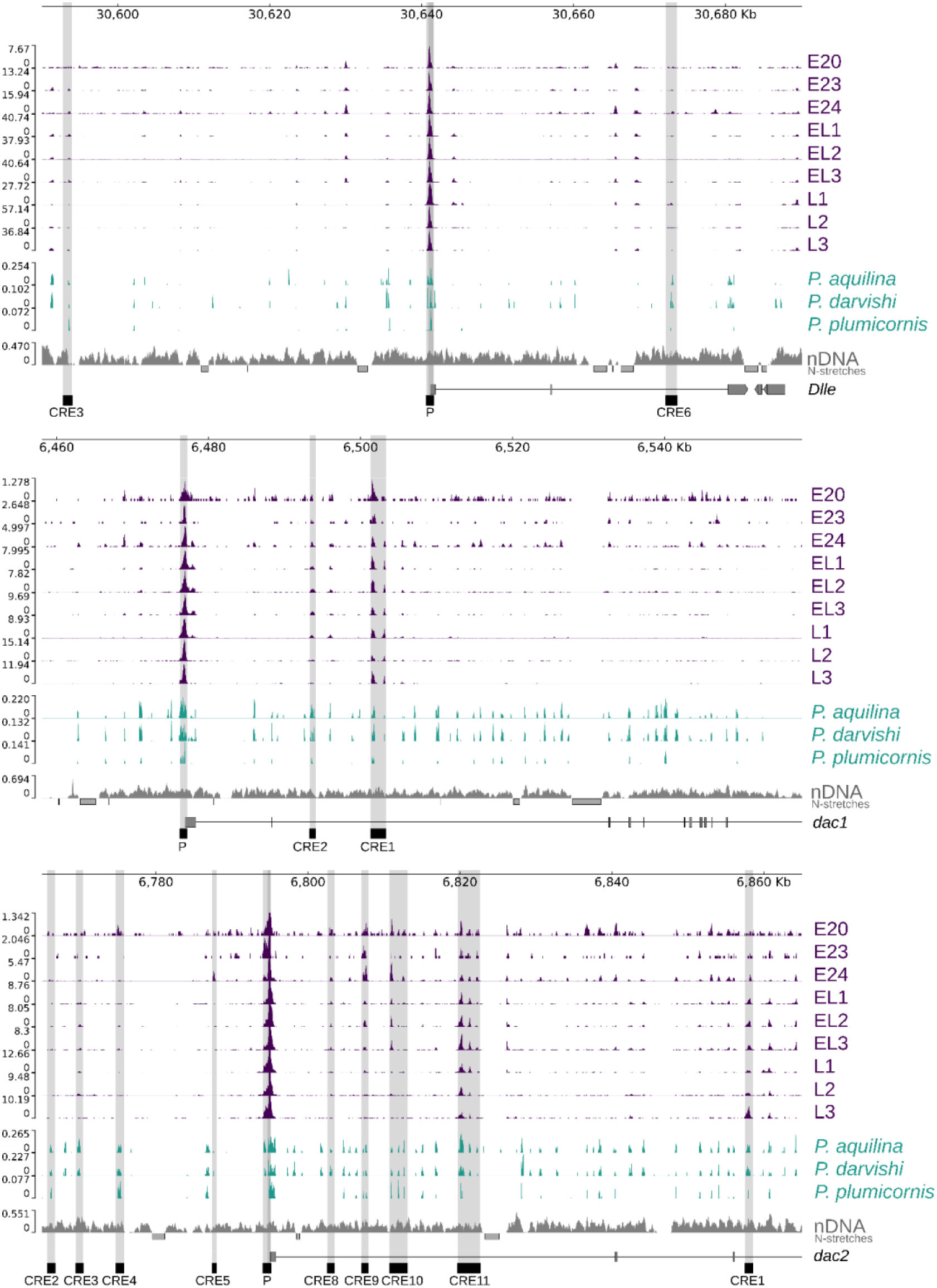
Putative CREs of *Parhyale* developmental genes tested using transgenic reporters. Genome browser plots for *Dll-e*, *dac1* and *dac2*, highlighting the promoter regions (P) and putative CREs that were tested (in grey). ATAC-seq, nDNA, and sequence conservation (*P. darvishi*, *P. aquilina*, and *P. plumicornis*) tracks were normalised by fragments per million and autoscaled to the highest peak in each region. For *dac2*, the y-axis maximum for whole embryo, embryo and adult leg tracks was halved to account for the large TSS peak, which otherwise masks the signal at putative CREs. Regions annotated as Ns are indicated by the N stretches track.

**Suppl. Figure 7.**
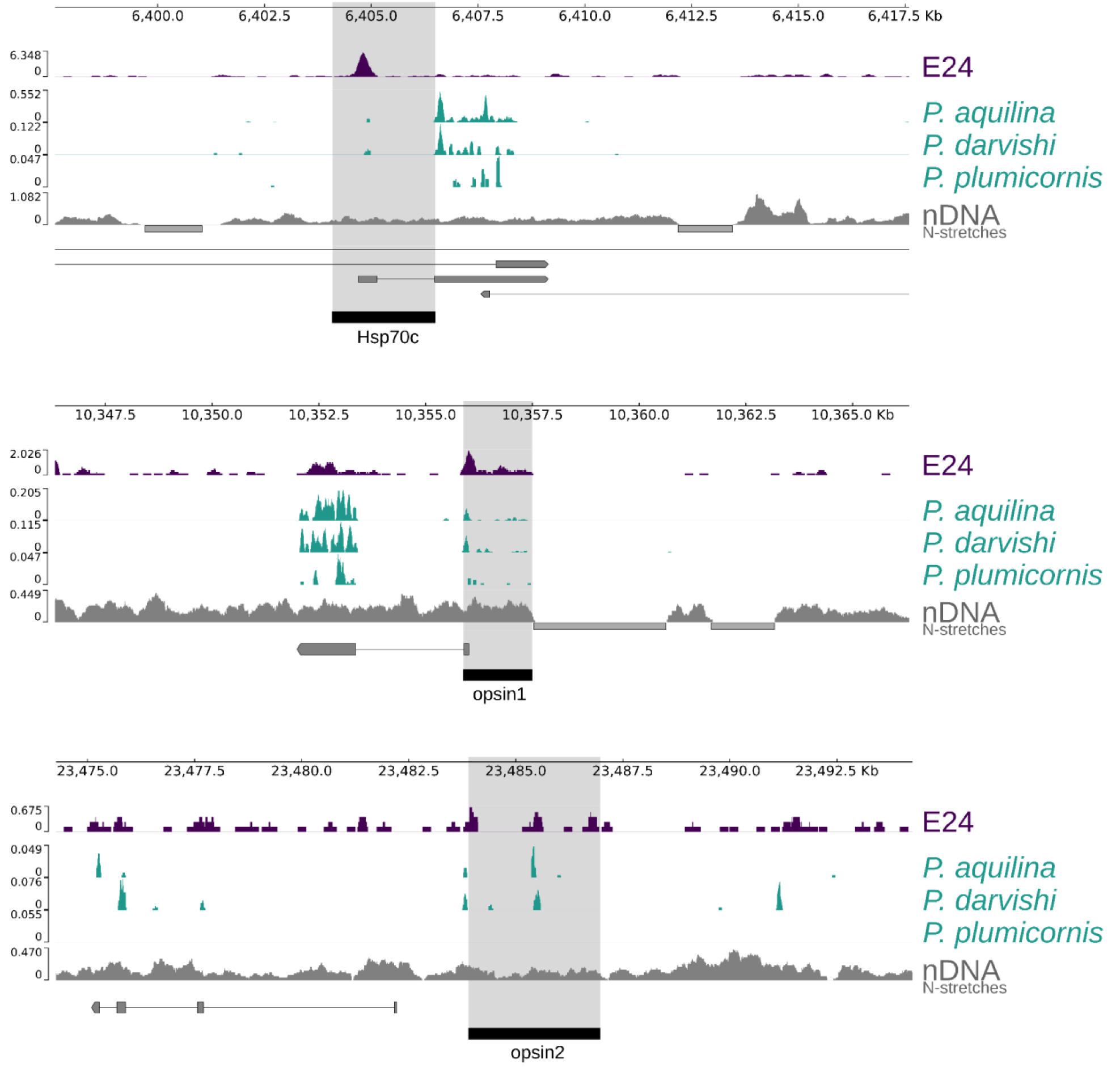
Chromatin accessibility and sequence conservation in previously identified *Parhyale* CREs. Genome browser plots showing the patterns of chromatin accessibility and cross-species sequence conservation in three genomic loci of *Parhyale hawaiensis*, that were previously shown to harbor active *cis-*regulatory elements: *PhMS* (*hsc70* locus, Pavlopoulos and Averof 2005), *PhOpsin1* and *PhOpsin2* (Ramos et al. 2019); the three elements are highlighted in grey. We observe ATAC-seq peaks and overlapping islands of sequence conservation among *Parhyale* species in all three fragments. Note that the same elements showed no significant sequence conservation when compared with homologous regions from the more distant species *Hyalella azteca* (Suppl. Figure 1).

**Supplementary Data Files** – available at https://zenodo.org/records/19020963

‣ Suppl. Data 1. Genome browser tracks and peaks for bulk ATAC-seq data
‣ Suppl. Data 2. Genome browser tracks for single-nuclei ATAC-seq data
‣ Suppl. Data 3. Genome browser tracks and islands of sequence conservation
‣ Suppl. Data 4. Sequence alignment for phylogeny
‣ Suppl. Data 5. DNA sequences and genome coordinates of all the tested CREs
‣ Suppl. Data 6. Microscopy data

## Notes

https://zenodo.org/records/19020963

### Competing Interest Statement

The authors have declared no competing interest.

### Summary of Updates

Minor revision following the suggestions of peer reviewers.

